# Development and evaluation of a decision prediction tool for the reduction of fungicide applications for the control of strawberry powdery mildew epidemics

**DOI:** 10.1101/2021.08.04.455115

**Authors:** Jolyon L. A. Dodgson, Bo Liu, Hannah J. Wileman, Euphemia S Mutasa-Gottgens, Avice M. Hall

## Abstract

Strawberry powdery mildew (*Podosphaera aphanis*) causes serious losses in UK crops, potentially reducing yields by as much as 70%. Consequently, conventional fungicide application programmes tend to recommend a prophylactic approach using insurance sprays, risking the development of fungicide insensitivity and requiring careful management relative to harvest periods to avoid residual fungicides on harvested fruit. This paper describes the development of a prediction system to guide the control of *P*. *aphanis* by the application of fungicides only when pathogen infection and disease progression are likely. The system was developed over a 15-year period on commercial farms starting with its establishment, validation and then deployment to strawberry growers. This involved three stages: 1. Identification and validation of parameters for inclusion in the prediction system (2004-2008). 2. Development of the prediction system in compact disc format (2009-2015). 3. Development and validation of the prediction system in a web-based format and cost-benefit analysis (2016-2020). The prediction system was based on the temporal accumulation of conditions (temperature and relative humidity) conducive to the development of *P. aphanis*, which sporulates at 144 accumulated disease-conducive hours. Sensitivity analysis was performed to refine the prediction system parameters. Field validation of the results demonstrated that to effectively control disease, the application of fungicides was best done between 125 and 144 accumulated hours of disease-conducive conditions. A cost-benefit analysis indicated that, by comparison with the number and timing of fungicide applications in conventional insurance disease control programmes, the prediction system enabled good disease control with significantly fewer fungicide applications (between one and four sprays less) (df=7, t=7.6, *p*=0.001) and reduced costs (savings between £35-£493/hectare) (df=7, t=4.0, *p*=0.01) for the growers.

## Introduction

The yield of the strawberry crop in the UK has more than trebled from the same area in the period 1995 to 2019, with the overall yield increasing from 41,600 tonnes in 1995 to 141,600 tonnes in 2019, giving a market value of £347.8 million [1]. This has been achieved through judicious use of short-day (June bearers) and day-neutral (everbearers) cultivars, polythene tunnels (S1 Fig), careful nutrition (fertigation), close planting (e.g. six plants per one meter coir bag), and careful management of the environmental conditions in the tunnels and grown in substrate [2]. Strawberry powdery mildew caused by *Podosphaera aphanis* ((Wallr.) Braun & Takamatsu) is a common disease worldwide [3], and it is the most serious epidemic disease threatening the strawberry crop grown in polythene tunnels in the UK (the majority of strawberry crops in the UK are now grown under polythene tunnels) [2]. Losses in crop yield can range from 20% to 70%; where a 20% loss at 2019 prices would have given a market value loss of £69.6 million [1, 2]. Cultivar choice by growers is influenced by a number of agronomic qualities, with resistance or tolerance to diseases being a low priority compared to aesthetic appeal, flavour and taste. As the potential loss from strawberry powdery mildew is great, growers generally resort to insurance (or prophylactic) spraying by applying a fungicide at one to two weekly intervals from April to September/October (this applies for both June bearers, which may have two plantings a year, and everbearers, with one planting a year), resulting in up to 24 fungicide applications per growing season and hence a higher incidence of fungicide residues compared to insecticide residues [4]. Insurance sprays cost money and can result in adverse environmental impacts, such as fungicide residues entering the soil or waterways due to spray drift or run-off [5].

Strawberry plants are supplied to growers by specialist propagators. In the UK, most strawberry crops are now grown in substrate (coir) on raised beds (S1 Fig a) or tabletops (S1 Fig d) or directly in the soil on raised beds. The crop is irrigated and fertilised via drippers inserted into the substrate (fertigation). There is a trend for annual planting to start in February, and there can be a very low incidence of *P. aphanis* on these plants when they are delivered to the grower by the propagator. Crops for early harvest are often covered with horticultural fleece to increase the temperature and prevent frost damage (S1 Fig b); this also increases the relative humidity under the fleece. This intensive production process has both extended the length of the UK growing season (harvest from late May to September/October) and increased the yield per hectare from 12.6 tonnes in 1999 to 29.7 tonnes in 2019 [2]. The use of polythene tunnels and other agronomic practices aim to provide optimal conditions for the growth of the crop, but it also provides ideal conditions for the growth of *P. aphanis*, risking epidemic development. Growers therefore need a reliable and efficient prediction system as a decision support system (DSS) to warn them when disease is most likely to occur and hence when to apply fungicides.

Prediction and forecasting systems have been developed and used on several crops since the 1970s [6–11]. However, uptake by growers has not been widespread or sustained, as the systems can be difficult to use and unreliable [6,7,11]. A prediction system is based on ‘formalized algorithms that assess disease risk factors to inform the need for crop protection’ [7]. It is a DSS that provides information (e.g. risk level of disease occurrence, timing of potential pest emergence etc.) to support the users (i.e. growers) in implementing the best strategies and practices in their disease and pest management [12, 13]. It should be emphasised here that DSSs only provide assistance, growers are the ones who make decisions [12]. DSSs can be developed based on the environmental conditions that follow the germination, growth and sporulation of the pathogen [6]. To be effective, a prediction DSS must be robust (i.e. the crop does not develop economically damaging disease during the season), simple to access, user-friendly, meet the needs of growers and help them to save money. The development of modern temperature and humidity sensors for use in crops, and the use of wireless web-based technology, offer a unique opportunity to develop a real-time localised DSS. The large number of insurance fungicide sprays currently recommended by crop advisors and used prophylactically each season by strawberry growers means that the development of a DSS based on reliable disease high risk prediction has the potential to reduce the number of fungicide applications. This offers growers financial incentives through savings from reduced fungicide sprays while maintaining disease control, with added environmental benefits. The rule-based prediction system developed and validated here applies to the asexual life cycle of *P. aphanis* (Fig 1), and we have successfully demonstrated its application as a DSS. Whilst there is some evidence of genetic variability in *P. aphanis* [14], there is no evidence of race specific pathotypes developing with respect to cultivars.

**Fig 1.**
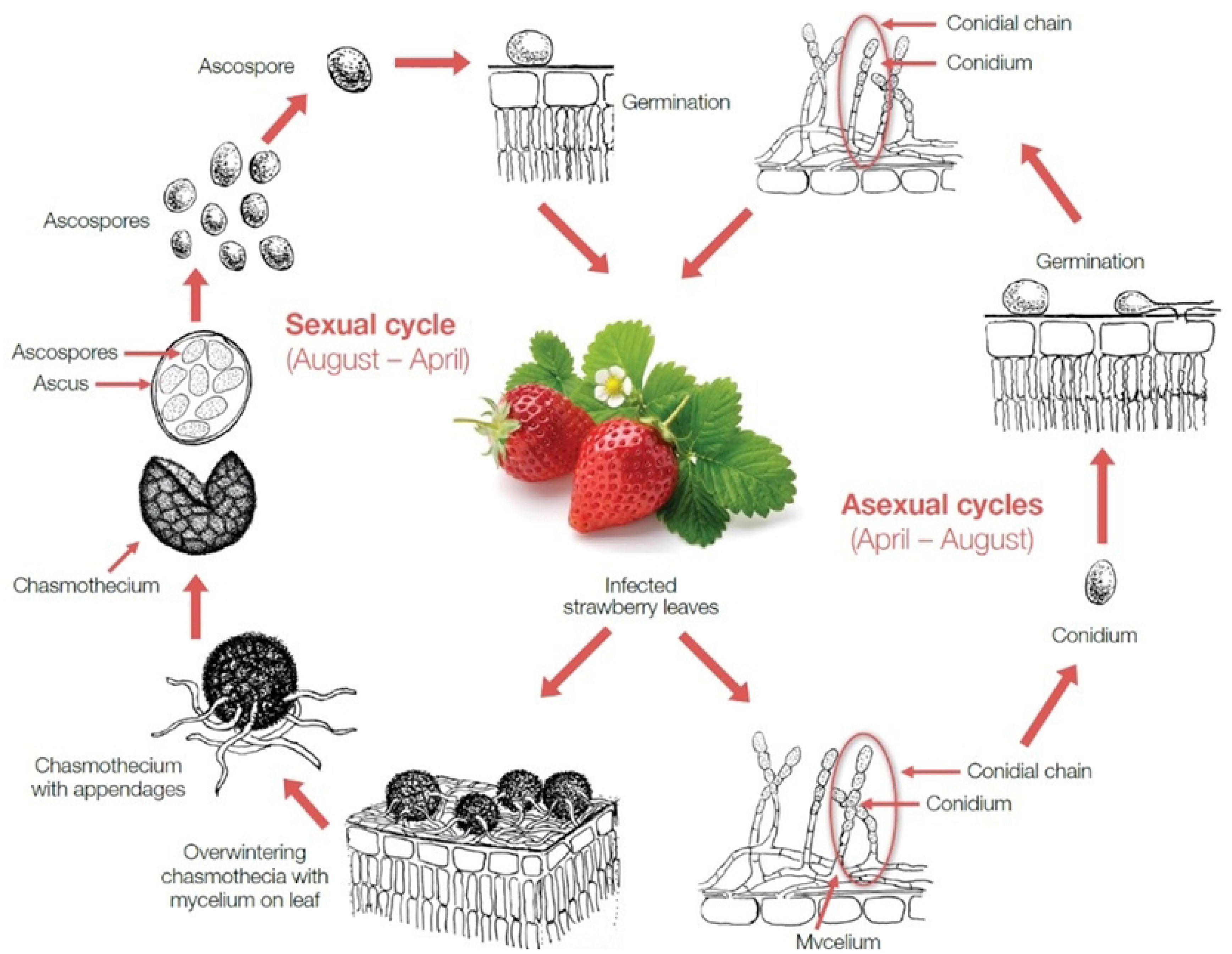
Sexual and asexual life cycles of *P. aphanis*. The sexual cycle occurs during the overwintering period when conditions are less conducive for pathogen growth. The asexual cycle occurs during the spring/summer period when the environmental conditions are suitable for growth, and this cycle is completed when there have been 144 accumulated hours of suitable conditions [1].

## Materials and methods

The development of the DSS included three stages (Table 1).

**Table 1.**
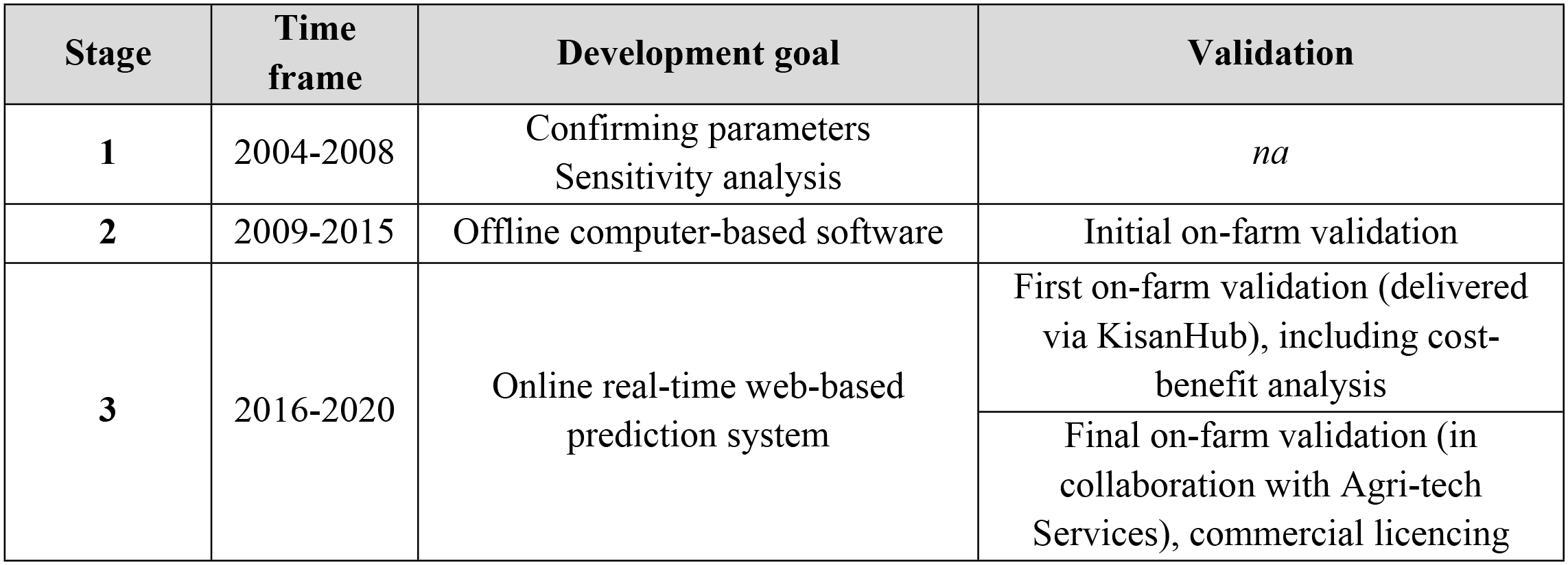
Development stages for the system for prediction of strawberry powdery mildew.

| Stage | Time frame | Development goal | Validation |
| --- | --- | --- | --- |
| 1 | 2004-2008 | Confirming parameters<br>Sensitivity analysis | *na* |
| 2 | 2009-2015 | Offline computer-based software | Initial on-farm validation |
| 3 | 2016-2020 | Online real-time web-based prediction system | First on-farm validation (delivered via KisanHub), including cost-benefit analysis |
|  |  |  | Final on-farm validation (in collaboration with Agri-tech Services), commercial licencing |

### Field sites used during development and validation of prediction system

The initial development of the strawberry powdery mildew prediction system (stage 1), involving parameter and sensitivity analysis, was done on four sites (Table 2), at commercial strawberry farms. Two sites had established strawberry plantings (cv. Elsanta) in their second year of growth, having remained in the ground over winter; these were near Mereworth (ME18) in 2004 and Wisbech (PE14) in 2005. Two sites were newly planted near Mereworth in 2005 (cv. Florence) and Wisbech in 2006 (cv. Elsanta). Within each field site, in-crop environmental conditions were recorded hourly using TinyTag Plus data loggers (Table 2) located within the crop canopy, with Tiny Tag Plus TGP-1500 for temperature and relative humidity and TinyTag Plus TGP-0903 for leaf wetness.

**Table 2.**
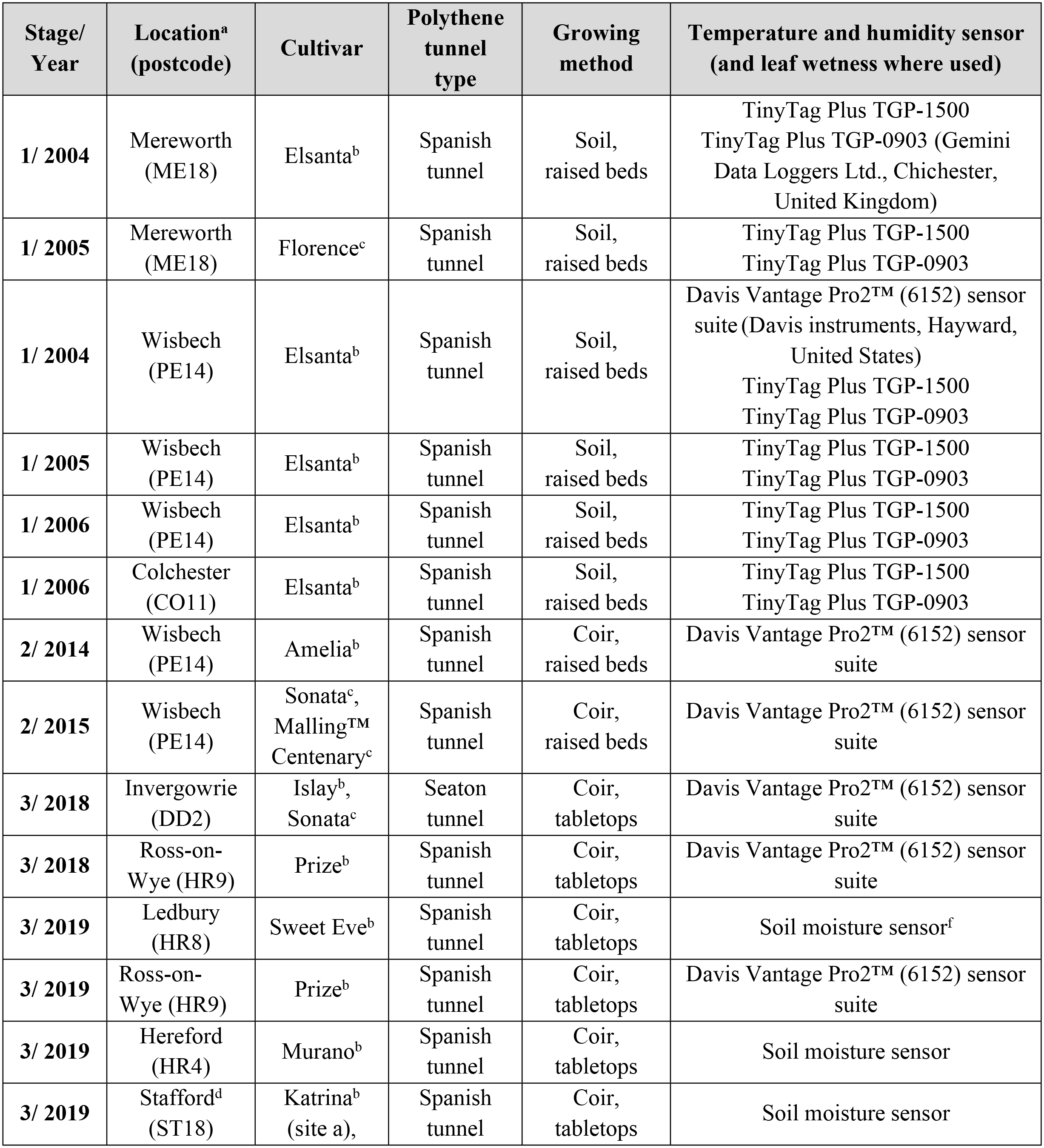
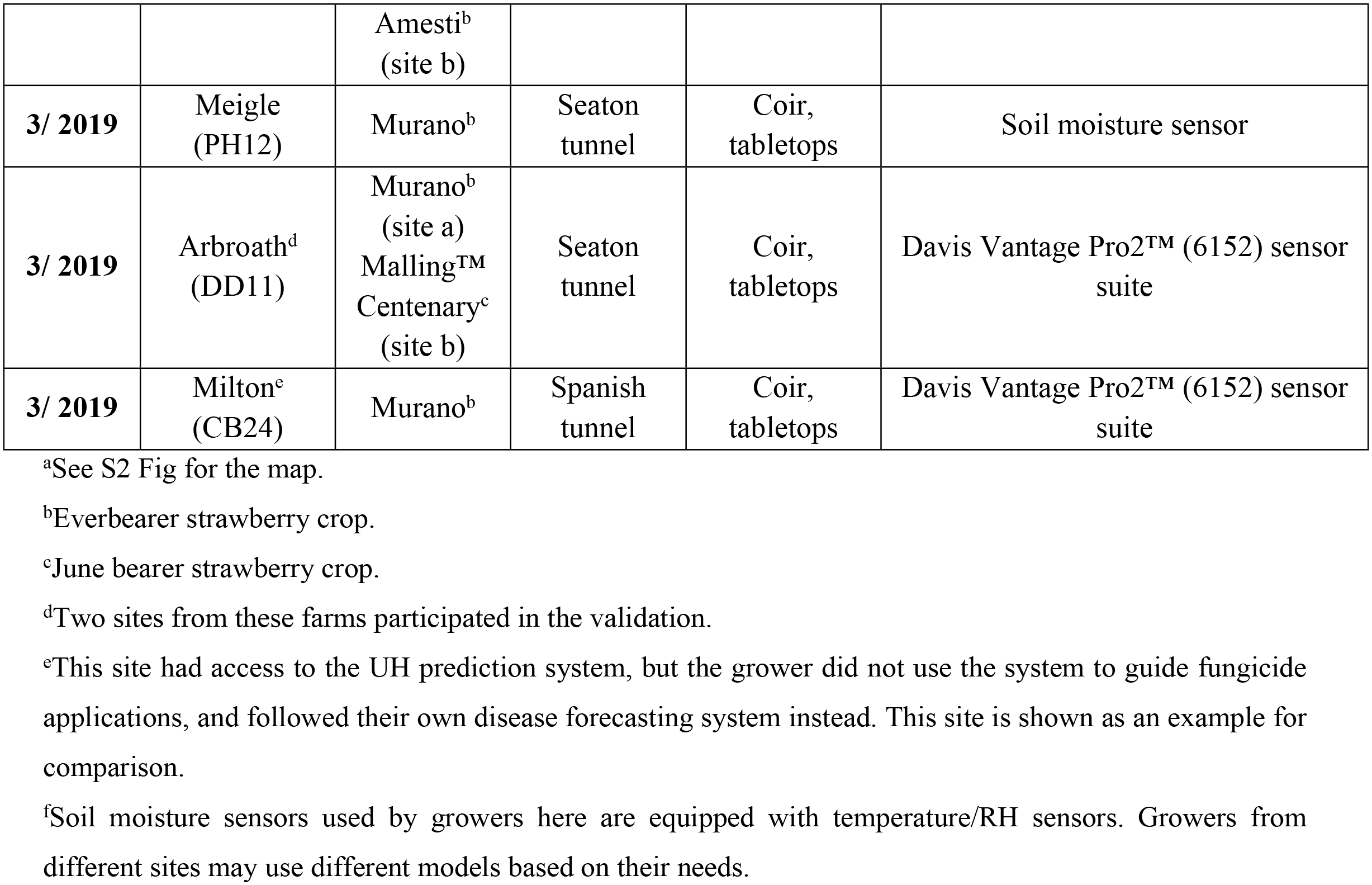
Project stage and year, site location, strawberry cultivar, growing methods and sensors used at all field sites at different stages in the development and validation of the strawberry powdery mildew prediction system.

| Stage/<br>Year | Location<sup>a</sup><br>(postcode) | Cultivar | Polythene<br>tunnel<br>type | Growing<br>method | Temperature and humidity sensor<br>(and leaf wetness where used) |
| --- | --- | --- | --- | --- | --- |
| 1/ 2004 | Mereworth<br>(ME18) | Elsanta<sup>b</sup> | Spanish<br>tunnel | Soil,<br>raised beds | TinyTag Plus TGP-1500<br>TinyTag Plus TGP-0903 (Gemini<br>Data Loggers Ltd., Chichester,<br>United Kingdom) |
| 1/ 2005 | Mereworth<br>(ME18) | Florence<sup>c</sup> | Spanish<br>tunnel | Soil,<br>raised beds | TinyTag Plus TGP-1500<br>TinyTag Plus TGP-0903 |
| 1/ 2004 | Wisbech<br>(PE14) | Elsanta<sup>b</sup> | Spanish<br>tunnel | Soil,<br>raised beds | Davis Vantage Pro2™ (6152) sensor<br>suite (Davis instruments, Hayward,<br>United States)<br>TinyTag Plus TGP-1500<br>TinyTag Plus TGP-0903 |
| 1/ 2005 | Wisbech<br>(PE14) | Elsanta<sup>b</sup> | Spanish<br>tunnel | Soil,<br>raised beds | TinyTag Plus TGP-1500<br>TinyTag Plus TGP-0903 |
| 1/ 2006 | Wisbech<br>(PE14) | Elsanta<sup>b</sup> | Spanish<br>tunnel | Soil,<br>raised beds | TinyTag Plus TGP-1500<br>TinyTag Plus TGP-0903 |
| 1/ 2006 | Colchester<br>(CO11) | Elsanta<sup>b</sup> | Spanish<br>tunnel | Soil,<br>raised beds | TinyTag Plus TGP-1500<br>TinyTag Plus TGP-0903 |
| 2/ 2014 | Wisbech<br>(PE14) | Amelia<sup>b</sup> | Spanish<br>tunnel | Coir,<br>raised beds | Davis Vantage Pro2™ (6152) sensor<br>suite |
| 2/ 2015 | Wisbech<br>(PE14) | Sonata<sup>c</sup>,<br>Malling™<br>Centenary<sup>c</sup> | Spanish<br>tunnel | Coir,<br>raised beds | Davis Vantage Pro2™ (6152) sensor<br>suite |
| 3/ 2018 | Invergowrie<br>(DD2) | Islay<sup>b</sup>,<br>Sonata<sup>c</sup> | Seaton<br>tunnel | Coir,<br>tabletops | Davis Vantage Pro2™ (6152) sensor<br>suite |
| 3/ 2018 | Ross-on-Wye (HR9) | Prize<sup>b</sup> | Spanish<br>tunnel | Coir,<br>tabletops | Davis Vantage Pro2™ (6152) sensor<br>suite |
| 3/ 2019 | Ledbury<br>(HR8) | Sweet Eve<sup>b</sup> | Spanish<br>tunnel | Coir,<br>tabletops | Soil moisture sensor<sup>f</sup> |
| 3/ 2019 | Ross-on-Wye (HR9) | Prize<sup>b</sup> | Spanish<br>tunnel | Coir,<br>tabletops | Davis Vantage Pro2™ (6152) sensor<br>suite |
| 3/ 2019 | Hereford<br>(HR4) | Murano<sup>b</sup> | Spanish<br>tunnel | Coir,<br>tabletops | Soil moisture sensor |
| 3/ 2019 | Stafford<sup>d</sup><br>(ST18) | Katrina<sup>b</sup><br>(site a), | Spanish<br>tunnel | Coir,<br>tabletops | Soil moisture sensor |
|  |  | Amesti<sup>b</sup><br>(site b) |  |  |  |
| **3/ 2019** | Meigle<br>(PH12) | Murano<sup>b</sup> | Seaton<br>tunnel | Coir,<br>tabletops | Soil moisture sensor |
| **3/ 2019** | Arbroath<sup>d</sup><br>(DD11) | Murano<sup>b</sup><br>(site a)<br>Malling<sup>TM</sup><br>Centenary<sup>c</sup><br>(site b) | Seaton<br>tunnel | Coir,<br>tabletops | Davis Vantage Pro2<sup>TM</sup> (6152) sensor<br>suite |
| **3/ 2019** | Milton<sup>e</sup><br>(CB24) | Murano<sup>b</sup> | Spanish<br>tunnel | Coir,<br>tabletops | Davis Vantage Pro2<sup>TM</sup> (6152) sensor<br>suite |

<sup>a</sup>See S2 Fig for the map.
<sup>b</sup>Everbearer strawberry crop.
<sup>c</sup>June bearer strawberry crop.
<sup>d</sup>Two sites from these farms participated in the validation.
<sup>e</sup>This site had access to the UH prediction system, but the grower did not use the system to guide fungicide applications, and followed their own disease forecasting system instead. This site is shown as an example for comparison.
<sup>f</sup>Soil moisture sensors used by growers here are equipped with temperature/RH sensors. Growers from different sites may use different models based on their needs.

In stage 2, the initial validation of the offline computer-based software was done at the Wisbech site (Table 2). One everbearer crop (cv. Amelia) was used in 2014 and two June bearer crops (cv. Sonata and Malling™ Centenary) were used in 2015. A Davis Vantage Pro2™ (6152) weather sensor suite was used to collect temperature and RH data.

In stage 3, two sites were used for the initial validation of the real-time web-based prediction system. One field site was situated near Invergowrie, Dundee (DD2), with an everbearer crop (cv. Islay) and two plantings of a June bearer crop (cv. Sonata) from March to June and June to October. Another site was located near Ross-on-Wye (HR9) growing an everbearer strawberry crop (cv. Prize) from March to October. The wide geographical area and representative range of cultivars used for validation was to enable the prediction system to be tested and subsequently used on any cultivar and throughout the UK.

The final on-farm validation (stage 3) (in collaboration with Agri-tech Services) took place on eight sites from seven farms in England and Scotland. Details of these sites are provided in Table 2.

### Assessment methods used for strawberry powdery mildew

In stage 1, strawberry crops were assessed at the field sites for the progression of the incidence of strawberry powdery mildew, caused by *P. aphanis.* Incidence (% plants affected) assessment was based on the presence of leaf upward cupping, visible fungal mycelium and/or red blotches on leaves (Fig 2). In spring (March-May) the fleece was removed from the strawberry beds and polythene covers were replaced onto the tunnel frames, having been removed during the winter to mitigate potential storm damage. Once this was completed, the regular assessment of plants began, until the symptoms of *P. aphanis* were fully developed throughout the untreated polythene tunnel (no applications of fungicides), so the incidence of strawberry powdery mildew was not inhibited by fungicides.

**Fig 2.**
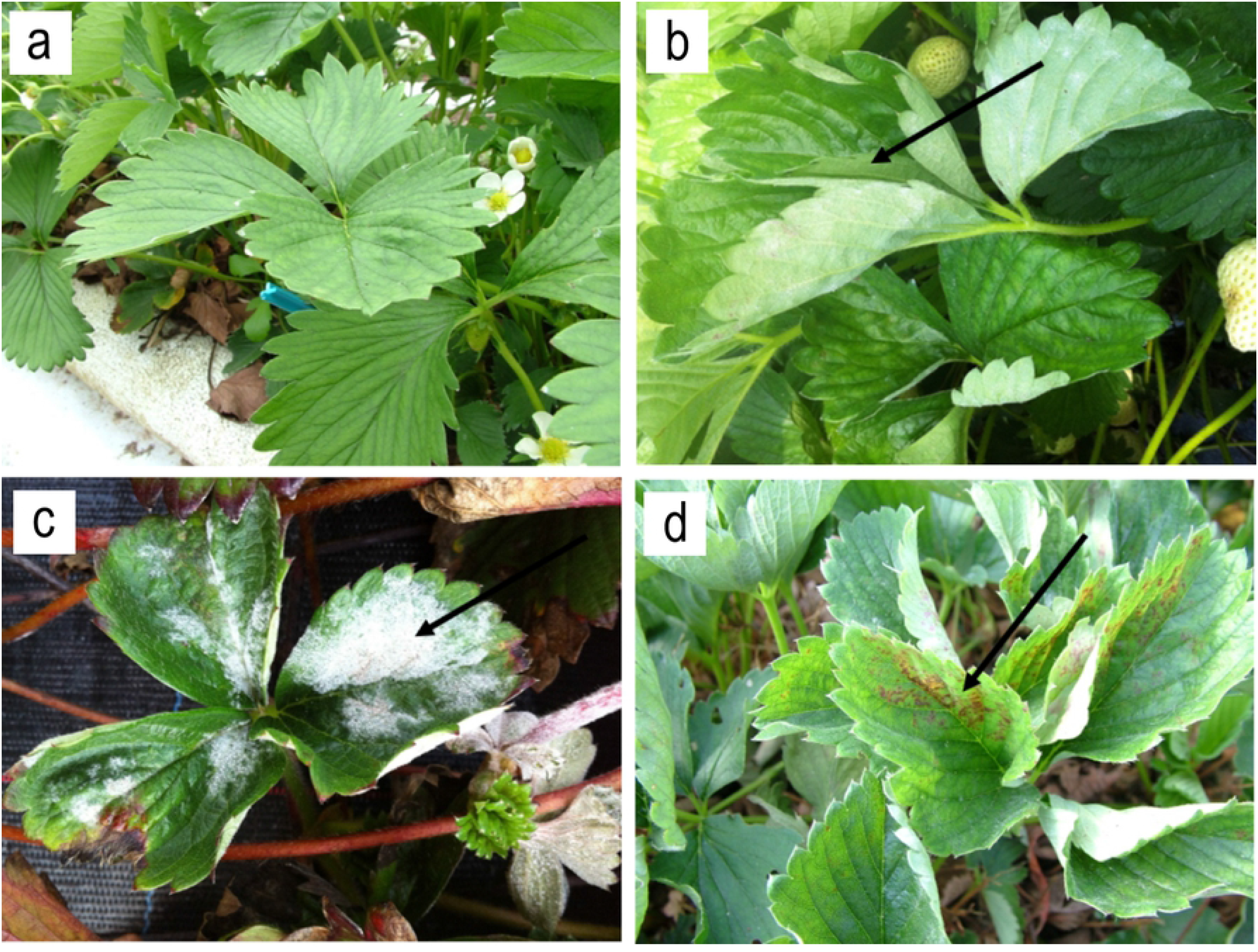
Symptoms of strawberry powdery mildew (arrows indicate the symptoms). (a) Healthy leaf on cultivar ‘Elsanta’; (b) cultivar ‘Amesti’ showing cupping; (c) cultivar ‘Amesti’ showing *P. aphanis* mycelium at 60% of leaf area affected, according to a strawberry powdery mildew disease assessment key [15]; (d) cultivar ‘Elsanta’ showing red blotches.

In stage 2, disease assessment was done by taking weekly strawberry leaf samples, from 01 July to 28 August in 2014 and from 01 July to 18 August in 2015. In 2014, the number of *P. aphanis* colonies on the leaf surface were counted and in 2015 the percentage leaf area covered by *P. aphanis* was assessed by using a new standard strawberry powdery mildew assessment key [15]. This updated assessment method was implemented due to it being a new industry standard that had been developed since the start of the work. Area Under the Disease Progress Curve (AUDPC) (The American Phytopathological Society, https://www.apsnet.org) was calculated to assess the level of *P. aphanis* development.

In stage 3, disease assessments were made at all farm sites by the growers. Disease control was considered to be good if no disease symptoms were observed, or if the disease level was very low and did not exceed commercially acceptable levels (i.e. no visible mycelium present on the fruits). On the other hand, if the disease level was considered by the growers to have exceeded commercially acceptable levels, the control was poor.

### Parameter and sensitivity analysis (stage 1)

#### Identification of environmental parameters that affected development of *P. aphanis*

The environmental parameters that were most likely to influence the asexual cycle of *P. aphanis* were identified from the published literature [16–22]. These environmental parameters (temperature, relative humidity and leaf wetness) were then used as the initial parameters for the strawberry powdery mildew prediction system (Table 3). The system was developed by computation using an Excel spreadsheet, into which the measurements of temperature, relative humidity and leaf wetness from sensors within crops grown in polythene tunnels were manually entered. The spreadsheet then calculated and identified when there had been a time period (one-hour increments) that was suitable for the growth and development of *P. aphanis*. The outputs of disease-conducive and non-conducive hours from this initial strawberry powdery mildew prediction system were then compared to the observed development of strawberry powdery mildew symptoms in a strawberry crop that had not been treated with fungicide at site PE14 (2006). The initial parameters were then modified so that the first predicted development of strawberry powdery mildew symptoms was synchronised with the timing of the symptoms observed in the crop. The modifications were a reduction in the minimum germination temperature, increase in the minimum growth temperature and changes to the elapsed time from asexual spore (conidia) germination to production of new spores of *P. aphanis*. These updated new field parameters (Table 3) were then used for the sensitivity analysis.

**Table 3.** Parameters derived from published laboratory-based methods (initial parameters^a^) and modified values after analysis of disease development data from commercial strawberry crop field sites (new field parameters).

| Pathogen development stage parameter | Initial parameters | New field parameters |
| --- | --- | --- |
| Minimum germination temperature (°C) | 17.5 | 15.5 |
| Maximum germination temperature (°C) | 30 | 30 |
| Minimum growth (and spore release) temperature (°C) | 16 | 18 |
| Maximum growth (and spore release) temperature (°C) | 30 | 30 |
| **Minimum relative humidity (%)** | 60 | 60 |
| **Minimum leaf wetness (%)<sup>b</sup>** | 95 | 95 |
| **No. of hours<sup>c</sup> for germination** | 6 | 6 |
| **No. of hours<sup>c</sup> for growth of elongating secondary hyphae to sporulation** | 138 | *na* |
| **No. of hours<sup>c</sup> for germination and growth of elongating secondary hyphae to sporulation in *established* field for first symptoms** | *na* | 6<sup>d</sup> + 78<sup>e</sup> |
| **No. of hours<sup>c</sup> for germination and growth of elongating secondary hyphae to sporulation in *established* field after first symptoms** | *na* | 6<sup>d</sup> + 138<sup>e</sup> |
| **No. of hours<sup>c</sup> for germination and growth of elongating secondary hyphae to sporulation in *new* field for all symptoms** | *na* | 6<sup>d</sup> + 138<sup>e</sup> |

Pathogen development stage from asexual spore (conidia) germination to production of new spores of *P. aphanis* is considered for the development of a strawberry powdery mildew prediction system.
<sup>a</sup>[16-22]
<sup>b</sup>This parameter was removed from the final prediction system following sensitivity analysis.
<sup>c</sup>Number of hours of suitable conditions (temperature, relative humidity and leaf wetness) for *P. aphanis*.
<sup>d</sup>Number of hours for germination.
<sup>e</sup>Number of hours for growth of elongating secondary hyphae to sporulation in an established/new field.

#### Sensitivity analysis

To determine which of the new field parameters had the greatest and smallest influences on the outputs of the strawberry powdery mildew prediction system, sensitivity analysis was done as previously described [23, 24]. In this case, the outputs of the strawberry powdery mildew prediction system were the number of disease-conducive hours, as determined by a rule-based system. Sensitivity analysis required keeping all the parameters constant, apart from the one being tested, which was altered in small increments of 0.5°C or 5% (RH) over a range of values (Table 4). This was to identify the parameter(s) that had the least impact on the prediction system, so that they could then be removed to simplify the rules of the prediction system. The data from these analyses were used to create the rules that determine the number of accumulated disease-conducive hours as the output of the prediction system.

**Table 4.** Sensitivity analysis of the new field parameters (Table 3) for determining rules of the strawberry powdery mildew prediction system.

| Prediction system parameter | Range of values | Increment of increase |
| --- | --- | --- |
| Minimum germination temperature (°C) | 13-25 | 0.5 |
| Maximum germination temperature (°C) | 25-35 | 0.5 |
| Minimum growth temperature (°C) | 13-25 | 0.5 |
| Maximum growth temperature (°C) | 25-35 | 0.5 |
| Minimum relative humidity (%) | 10-100 | 5 |
| Minimum leaf wetness (%) | 60-100 | 5 |

### Comparison of prediction system outputs to commercial insurance spray programmes (stage 1)

At the end of a given growing season, the number of high-risk periods when a strawberry grower was advised to use a control product (e.g. fungicide) for *P. aphanis* according to the prediction system was compared to the actual number of fungicide applications made according to the commercial routine spray programmes. To achieve this, environmental parameters were recorded hourly with TinyTag data loggers and downloaded to a computer for the duration of the growing season to provide input for the prediction system. The actual number of insurance applications made to crops of strawberries were obtained from the growers. At the field sites from stage 1 (Table 2), the growers managed disease control according to accepted commercial practice which meant that they applied a fungicide when, based on experience, they perceived the risk of *P. aphanis* to be high. By default, the strawberry growers followed a customary routine spray programme and applied what they considered to be insurance sprays [25] (spraying every 7-14 days regardless of disease severity) to mitigate the potential for *P. aphanis* to develop in their polythene tunnels.

### Development and validation of the prediction system as a decision support tool (stages 2-3)

The development and validation of the rule-based prediction system to support on-farm decision making involved two stages: the use of computer-based software with off-line infrastructure (stage 2) and the development of an online, fully connected real-time web-based system (stage 3). A summary of the validation process is presented in Table 5, with comparisons between these two stages.

**Table 5.** Development and comparison of the prediction system over stage 2 (offline computer-based software) and stage 3 (online real-time web-based system).

| Development stage | Period | Collection of weather data | Data transmission | Viewing of prediction graph | Validation period & method | Validation assessment | Notes |
| --- | --- | --- | --- | --- | --- | --- | --- |
| Stage 2 | 2009-2015 | On-farm weather station located within the polythene tunnel (Table 2) | Data were downloaded and input into computer-based Davis WeatherLink™ software | On computer, not real-time | 2014-2015, one farm at site PE14<br>Three treatments:<br>- untreated control<br>- prediction system (applying fungicide according to the prediction system)<br>- commercial practice (applying fungicide under normal farm insurance practice) | Weekly leaf samples were assessed in the laboratory at University of Hertfordshire (UH) for numbers of *P. aphanis* colonies (2014) or % leaf area covered by *P. aphanis* colonies (2015)<br>AUDPC values were calculated. | Entire process was manually operated by growers, not real time, time-consuming, not user-friendly |
| Stage 3 | 2016-2018 (Initial validation) | Davis wireless sensors located within the polythene tunnel (Table 2) | Data were transmitted via Davis WeatherLink™ online hourly | Online, real-time | 2018, two participating farms (England)<br>On-farm disease assessment by growers and cost-benefit analysis were done. | Disease assessment made by growers in the field on the standard of commercially acceptable levels<br>All season fungicide data and costings gathered by researchers at UH, number of sprays, mode of action used etc. compared between prediction and insurance programmes | Entire process was done online, graph displayed in real-time, time-saving<br>Presentation of accumulated hours confusing |
|  | 2019-2020 (final validation) | Davis or SMS wireless sensors located within the polythene tunnel (Table 2) | Data were transmitted via Davis WeatherLink™ online hourly | Online, real-time | Seven participating farms (England and Scotland)<br>Criteria:<br>- needs to provide commercially acceptable disease control<br>- needs to work at different geographical locations<br>- needs to work on different cultivars<br>- needs to work irrespective of cultivation methods<br>On-farm disease assessment by growers and cost-benefit analysis were done |  | Presentation of the graph improved, easy to read, the system was user-friendly |

#### Offline computer-based software (stage 2)

The rule-based prediction system in the form of a computer program was then developed by Prof Xiangming Xu (NIAB-EMR) [26], using the parameters identified from previous years (Table 4), and this was used in stage 2 on the farm at the site PE14 (Table 5). Growers were able to obtain the weather data from the weather station and used the program to calculate the number of accumulated disease conducive hours, hence to measure the level of disease risk. The efficacy of the prediction system in disease control (achieving commercially good disease control with reduced fungicide sprays) was assessed at the end of stage 2.

#### Online real-time web-based prediction system (stage 3)

##### Proof of concept

With the availability of the internet and wifi-enabled weather stations, sensors and smart devices, the rule-based prediction system was transferred to a web platform for real-time data acquisition and analysis. In stage 3, the system was delivered initially via the KisanHub platform (https://www.kisanhub.com/platform/) between 2016 and 2018, using the parameters for temperature and relative humidity developed and validated previously (Table 5). By viewing the prediction system output directly on-line, the grower could track the accumulation of disease-conducive hours in real time. Initial validations of the real-time prediction system were done on commercial farms (Table 2) and the usability with respect to data visualisation and user dashboard on the platform was assessed.

##### Final validation

Nine sites from seven farms took part in the final validation of the prediction system in 2019 (in collaboration with Agri-tech Services) (Tables 2 and 5). Fungicide data for the growing season, costings and disease assessment results were obtained for the nine sites from the participating farms. On receipt of the results from the growers, fungicide spray schedules were analysed for the number of sprays, the active ingredients (mode of action) used, spray intervals and the number of disease-conducive hours that had accumulated when a fungicide was applied. An objective evaluation of the use of the system was then made. Good use of the prediction system was when a fungicide application was made between 115 and 144 accumulated hours of disease-conducive conditions; the system was considered not to have been used to its full potential if fungicide applications were made below 115 hours; and the system was considered not to have been followed if a fungicide application was made at 50 accumulated hours or below.

##### Cost-benefit analysis

A cost-benefit analysis was done by calculating the price per hectare of each fungicide. The sum of the cost of all fungicide sprays was determined to give the total fungicide cost per hectare for the season. An indicative labour cost for a single fungicide application was given by growers and multiplied by the total number of sprays. The total cost of fungicides per hectare plus the total labour cost gave a total cost for fungicide applications per hectare for the season. The total cost of fungicide applications calculated when guided by the prediction system was compared to the total cost of fungicide applications when following a commercial (insurance) spray programme. The average cost of a single fungicide application per hectare (fungicide plus labour) was calculated and multiplied by the number of sprays applied when following an insurance spray programme, thereby giving an estimated cost for this spray programme (control). Data normality was determined using the Kolmogorov-Smimov Test [27] and paired T-tests were used to determine significant differences between the number of fungicide sprays and cost of fungicides and labour between the prediction system and insurance spray programmes.

#### Visualisation of prediction system output and workflow for the grower (stages 2-3)

The prediction system used a ‘traffic light’ colour scheme indication to represent the progression of accumulated hours of conditions conducive to *P. aphanis* development, visualised as an ascending line graph. Green indicated a low risk of *P. aphanis* sporulation. When 115 hours had accumulated, the line changed to amber, advising that the grower should prepare to apply a fungicide. When 125 accumulated hours were reached, the line changed to red, indicating a high risk of conidial sporulation and that a fungicide spray should be applied. A fungicide should normally be applied before the elapsed time reaches 144 accumulated hours, to prevent *P. aphanis* sporulation. When a fungicide application was made, the growers recorded this manually in the software and reset the system to zero accumulated hours; subsequent hours were then accumulated only when the temperature (15-30°C) and relative humidity (>60%) criteria were both met; such that the conditions within the polythene tunnel were conducive for *P. aphanis* development.

## Results

### Parameter and sensitivity analysis (stage 1)

#### Assessment of *P. aphanis*

The development of the first symptoms of *P. aphanis* infections was scored in four separate polyethene tunnels that were at two separate commercial sites, starting in spring and summer (March-July), when the tunnel frames were covered with polythene at the start of the growing season. Similar results were observed at all sites and Fig 3 shows representative results from the polythene tunnel at site ME18, where an epidemic occurred, with a lag phase, exponential phase and stationary phase of disease development. This also shows the accumulation of hours where the conditions were predicted to be suitable for the growth of *P. aphanis* when using the initial parameters (Table 3). If the initial parameters had provided a prediction that was a good fit to disease development in the crop, then the accumulation of disease-conducive conditions should have reached 100% just as the disease development was at the start of the exponential phase. However, this did not happen, because strawberry powdery mildew symptoms were observed before disease-conducive hours were predicted by the system using the initial parameters (Fig 3, line with open squares). Therefore, field observations were used to modify the initial parameters to give the new parameters (Table 3). These new parameters were then used to calculate the accumulation of disease-conducive conditions for the development of *P. aphanis* in the crop. In Fig 3, the line representing the new parameters (solid squares) reached 100% just as the first visible symptoms of strawberry powdery mildew were observed on the strawberry plants. As the prediction system is based on only those hours that are suitable for the initiation of germination and growth of *P. aphanis*, the prediction system must necessarily depict the accumulation of disease-conducive hours as defined by the new parameters.

**Fig 3.**
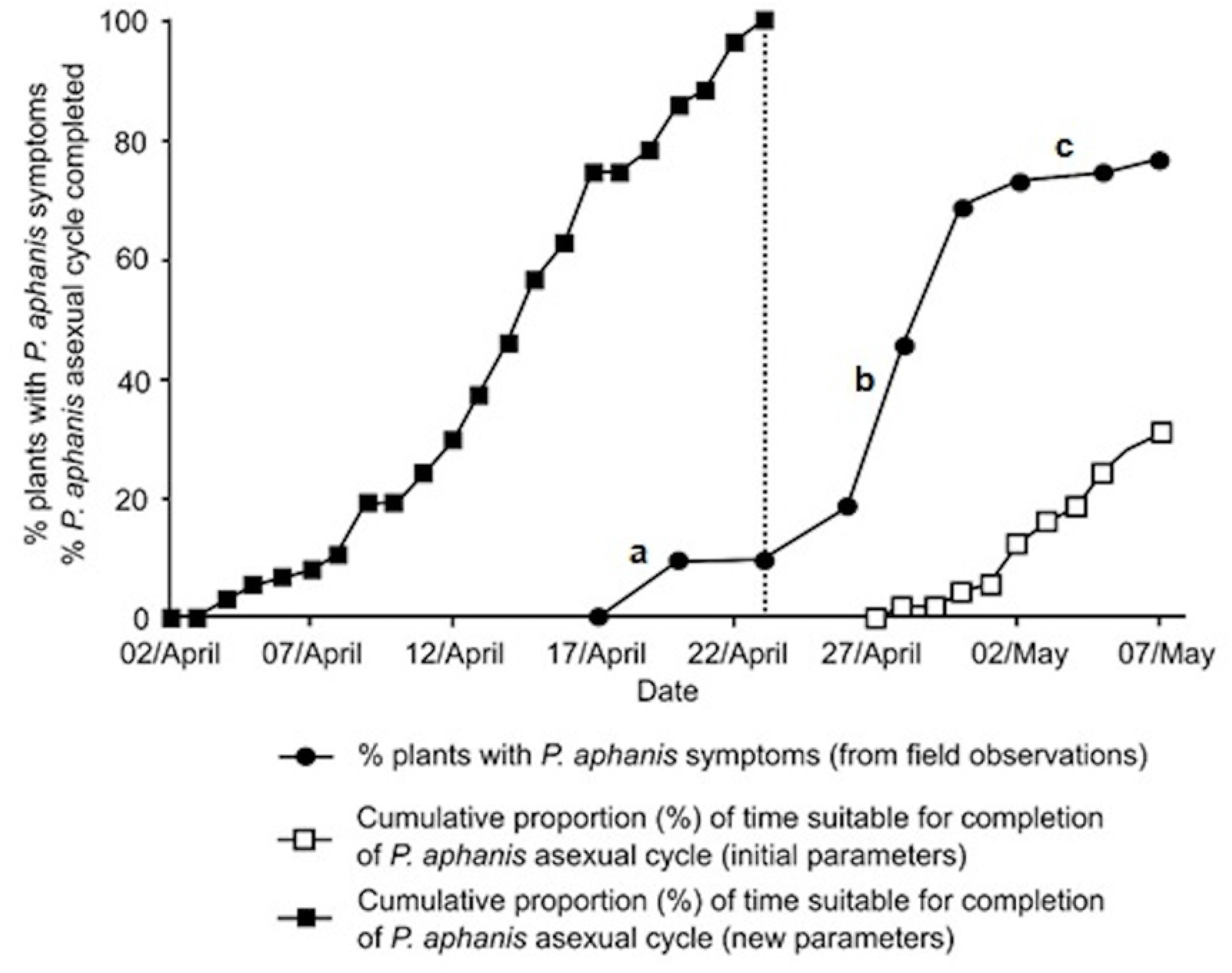
Percentage of strawberry leaves showing symptoms of *P. aphanis* infection in the polytunnel at field site ME18 (stage 1). The epidemic followed a disease progress curve with (a) a lag phase, (b) an exponential phase and (c) a stationary phase. 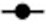 indicates % plants with *P. aphanis* symptoms observed in the field. The cumulative percentage of disease-conducive hours for the progression of *P. aphanis* asexual cycle (i.e. the time it would take for a conidium to germinate, colonise and produce more conidia, which is usually 144 hours of suitable conditions) was calculated when using the initial and new parameters (Table 3) for the strawberry powdery mildew prediction system. The initial parameters did not accurately predict the start of the epidemic; the disease progress (indicated 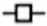) was in the exponential phase and symptoms had already appeared when the cumulative percentage was starting to increase. Therefore, using the initial parameters had not accurately predicted when to apply a fungicide spray. However, using the new parameters, the predicted cumulative percentage (indicated by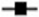) had reached 100% as the symptoms were starting to appear. Administering a fungicide application at 100% (indicated by dotted line) would coincide with the disease progress in the lag phase of the epidemic, aiding in reducing the initial inoculum [28].

#### Identification of environmental parameters that affected development of *P. aphanis*

The environmental parameters (i.e. temperature, relative humidity and leaf wetness required for germination, growth and sporulation of *P. aphanis*) are shown in Table 3. These parameters affected the development of *P. aphanis* by influencing the rate at which the pathogen germinated and grew, and therefore the amount of time needed for the pathogen to produce more conidia. Under laboratory conditions, it was identified that 144 hours of conducive conditions were required for the pathogen to germinate, grow and produce new conidia [22]. The information in Fig 4 demonstrates what happens during those 144 hours of suitable conditions. It should be noted that if the conditions are not suitable then the life cycle does not progress, but it also does not regress, and instead arrests, so that once conducive conditions are present again, the life cycle will resume from the stage it had reached previously. It is this potential for the resumption and continuation of development that the strawberry powdery mildew prediction system aims to disrupt and stop, by recommending precisely timed applications of fungicides while also giving growers the confidence not to apply insurance sprays when there have not been enough disease-conducive hours accumulated.

**Fig 4.**
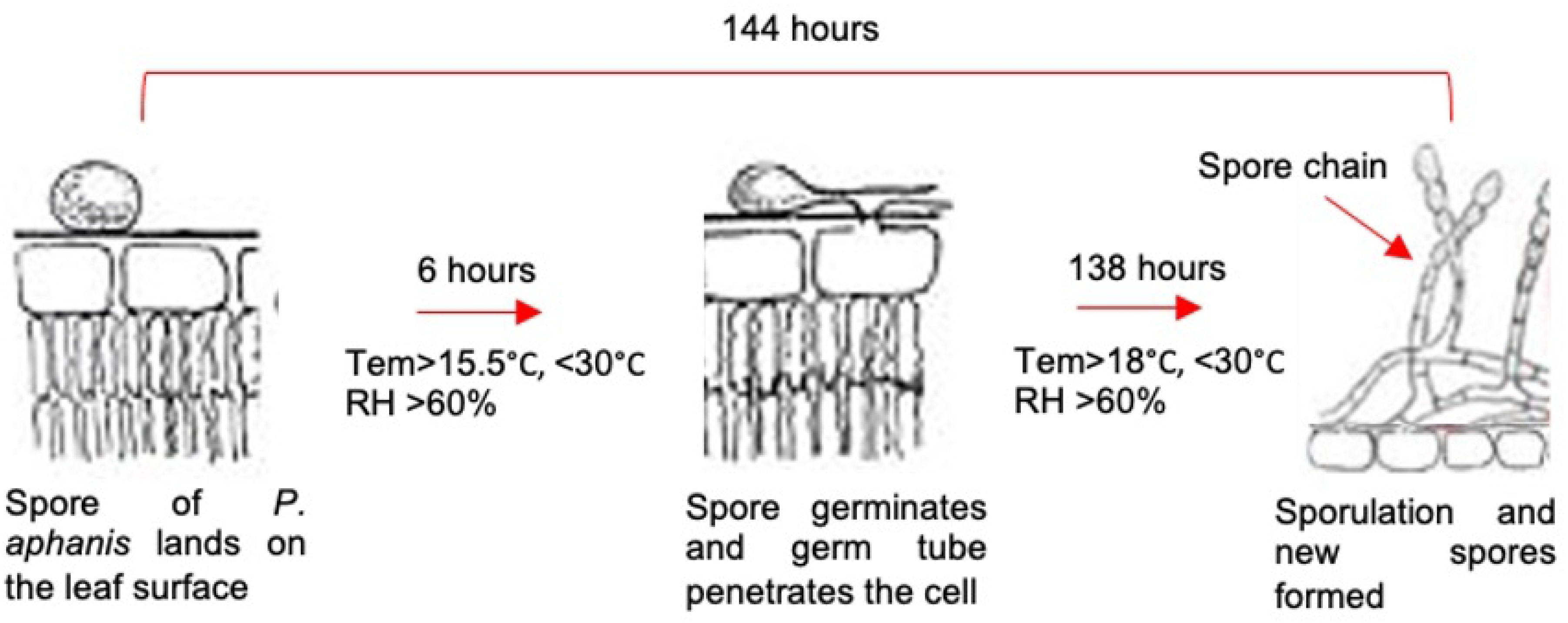
Diagram of the asexual life cycle of *Podosphaera aphanis* showing the parameters governing the rule-based prediction system. The system accumulates the number of hours when the conditions have been suitable for the development of *P. aphanis* (i.e. disease-conducive hours), assuming that it is not the first disease cycle in an established field. For *P. aphanis*, the conditions are temperature >15.5°C and <30°C (15.5°C is the minimum temperature for spore germination, whereas 18°C is the minimum temperature for sporulation), with relative humidity >60%. The asexual cycle starts with the germination of the conidia (c. six hours), followed by the growth of the pathogen (i.e. sporulation, c. 138 hours). Then once the number of suitable hours (144 hours) has been reached, the system recommends that the grower apply a control product for *P. aphanis*, after which the system resets itself and begins again. (Modified from Jin 2016.)

#### Sensitivity analysis

Once the new parameters (Table 3) had been shown to correspond well to the observed development of *P. aphanis* in the crop (Fig 3), a sensitivity analysis was done. This determined which of the parameters had the greatest and smallest influences on the total number of predicted completed asexual life cycles (referred to as high-risk periods) of *P. aphanis* in the crop. As shown in Table 6, leaf wetness had very little effect on the number of high-risk periods, when considered as a parameter on its own or in combination with each of the other parameters. The parameter that had the greatest influence on the number of high-risk periods was temperature. Following the sensitivity analysis, the parameters were refined again by removing leaf wetness from the prediction system, since it had a limited impact on the output.

**Table 6.** Results of sensitivity analysis on new parameters presented in Table 3.

| Prediction system parameter | Maximum number of times 100% of required disease-conductive hours were accumulated | Minimum number of times 100% of required disease-conductive hours were accumulated | Difference in number of times 100% of required disease-conductive hours were accumulated<sup>a</sup> |
| --- | --- | --- | --- |
| Minimum germination temperature with leaf wetness | 11 | 5 | 6 |
| Minimum germination temperature without leaf wetness | 11 | 7 | 4 |
| Maximum germination temperature with leaf wetness | 11 | 11 | 0 |
| Maximum germination temperature without leaf wetness | 11 | 11 | 0 |
| Minimum growth temperature with leaf wetness | 22 | 4 | 18 |
| Minimum growth temperature without leaf wetness | 23 | 4 | 19 |
| Maximum growth temperature with leaf wetness | 12 | 8 | 4 |
| Maximum growth temperature without leaf wetness | 12 | 8 | 4 |
| RH with leaf wetness | 11 | 4 | 7 |
| RH without leaf wetness | 11 | 5 | 6 |
| Leaf wetness | 11 | 10 | 1 |

<sup>a</sup>The difference in the number of times that 100% of the required disease-conductive hours for growth and conidial development elapsed indicates which parameters had the greatest and least impacts on the strawberry powdery mildew system; the prediction system parameter(s) that had the least impact could be removed to simplify the prediction system.

#### Comparison of prediction system outputs to commercial insurance spray programmes (stage 1)

To determine whether there were more fungicide applications being applied than were necessary, the number of predicted high-risk periods was compared to the actual number of insurance fungicide applications that growers used to control *P. aphanis*. The number of grower applications at site PE14 in stage 1 (2004 only) was 10 with five of those being applied during the period when the crop was being harvested, while the prediction system would have recommended eight applications with only three during the harvest period. At site CO11 in stage 1 (2006 only), the grower applied eight applications with two during the harvest period when the prediction system would have recommended five applications with two during the harvest period. At both sites, the growers were applying insurance sprays that would not have been done had they been guided by the prediction system. The disease-conducive hours took longer to develop to a high-risk period than estimated by the growers without reference to the prediction system.

The initial phase of developing the strawberry powdery mildew prediction system, using Excel, was done to demonstrate that it had the potential to reduce the number of applications of fungicides to control *P. aphanis*. Once this had been done, it was possible to relate the prediction system with stages of the asexual life cycle of *P. aphanis* to visualise the steps in the process (Fig 4).

### Validation of the prediction system (stages 2-3)

Results of the validation of the prediction system over stage 2 (offline computer-based) and stage 3 (online real-time web-based) are presented in Tables 7 and 8.

**Table 7.**
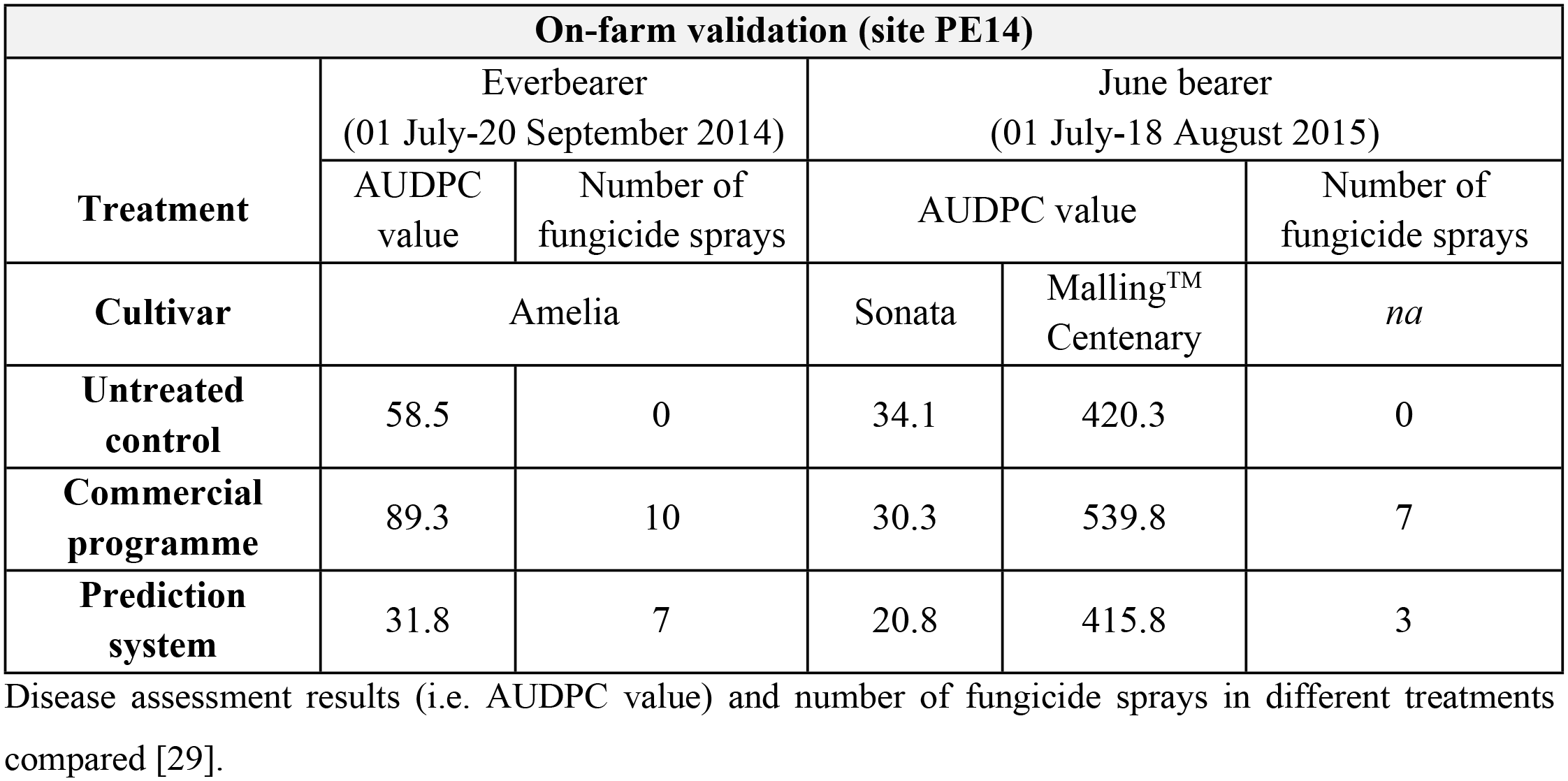
Validation results of the prediction system from offline computer-based software (stage 2).

**Table 8.** Validation results of the prediction system from the online real-time web-based system (stage 3).

| Initial validation |  |  |  |  |  |  |
| --- | --- | --- | --- | --- | --- | --- |
| Validation site | Site DD2<br>(Everbearer) |  | Site DD2<br>(June bearer) |  | Site HR9<br>(Everbearer) |  |
| Treatment | Number of fungicide sprays | Total cost £/hectare (fungicide +labour) | Number of fungicide sprays | Total cost £/hectare (fungicide +labour) | Number of fungicide sprays | Total cost £/hectare (fungicide +labour) |
| Commercial programme | 13 | 1,194.60 | 11 | 1,029.44 | 23 | 1658.98 |
| Prediction system | 10 | 918.92 | 8 | 748.68 | 19 | 1,442.65 |
| Saved | 3 | 275.68 | 3 | 280.76 | 4 | 216.33 |
| Grower disease assessment | Good control |  | Good control |  | Good control |  |

This table shows the number of fungicide sprays and total cost (fungicide + labour) compared between commercial and prediction programmes at three sites.

#### Computer-based software

The validation results in stage 2 showed that crops from the prediction system field plots had less disease (i.e. smaller AUDPC values) compared to those from the insurance spray programme (commercial) plots and untreated control (Table 7). Growers reported a low incidence of disease in the crop over the duration of the season; the prediction system gave good control (commercially acceptable) of strawberry powdery mildew, as an epidemic did not develop.

Disease assessment results (i.e. AUDPC value) and number of fungicide sprays in different treatments compared [29]. The prediction plot in 2014 received seven sprays compared to 10 in the commercial practice plot (Table 7). In 2015, the prediction plot was sprayed only three times using two fungicide active ingredients, compared to seven sprays with a total use of eight active ingredients in the commercial practice plot. The prediction system offered good control with fewer sprays compared to commercial plots, which also had higher levels of disease even though they were sprayed more frequently. Manual downloading of the data from the Davis Pro2^TM^ sensors via the Weatherlink™ software and then uploading it onto the prediction system software was required several times per week to obtain an accurate prediction of the high-risk periods, and this was time-consuming for the grower. Importantly, it has been successfully demonstrated that the prediction system had potential for commercial use if the data transfer and integration could be made less cumbersome, making the process more user-friendly.

#### Real-time web-based software

The initial validation in stage 3 showed that responding to the monitoring of the disease-conducive hours using the prediction system, gave longer intervals (>14 days) between fungicide applications at the start of the season, enabling the grower to save sprays. Overall, cost savings averaging £250 ha^-1^ were achieved at the sites where growers followed the prediction system (Table 8).

In the final validation, the prediction system was used on eight sites (six farms) resulting in varying levels of success, with a control site (CB24) where it was not used to support the application of fungicides but instead, the system collected the disease conducive hours in the polythene tunnels and the grower recorded when fungicides were applied using their insurance spray programme (Table 9). Some growers used the prediction system well (e.g. site HR8), whereas some did not use it to its full potential as indicated by the reduction in spray applications as well as the overall cost savings, ranging from £35/ha up to £493/ha. The results showed a significant difference (*p*<0.05) for both the number of sprays (df=7, t=7.6, *p*=0.001) and overall costs (df=7, t=4.0, *p*=0.01) between the prediction system and insurance spray programme. Every site, except CB24, was able to save at least one spray application by following the prediction system, and the most efficient (HR8), was able to reduce applications by as much as 50% compared to their insurance programme. The number of insurance sprays more or less correlated with the level of risk perception by the grower and data clearly showed that a large number of spray applications (e.g. sites CB24, ST18 and HR9) do not necessarily result in good disease control.

**Table 9.**
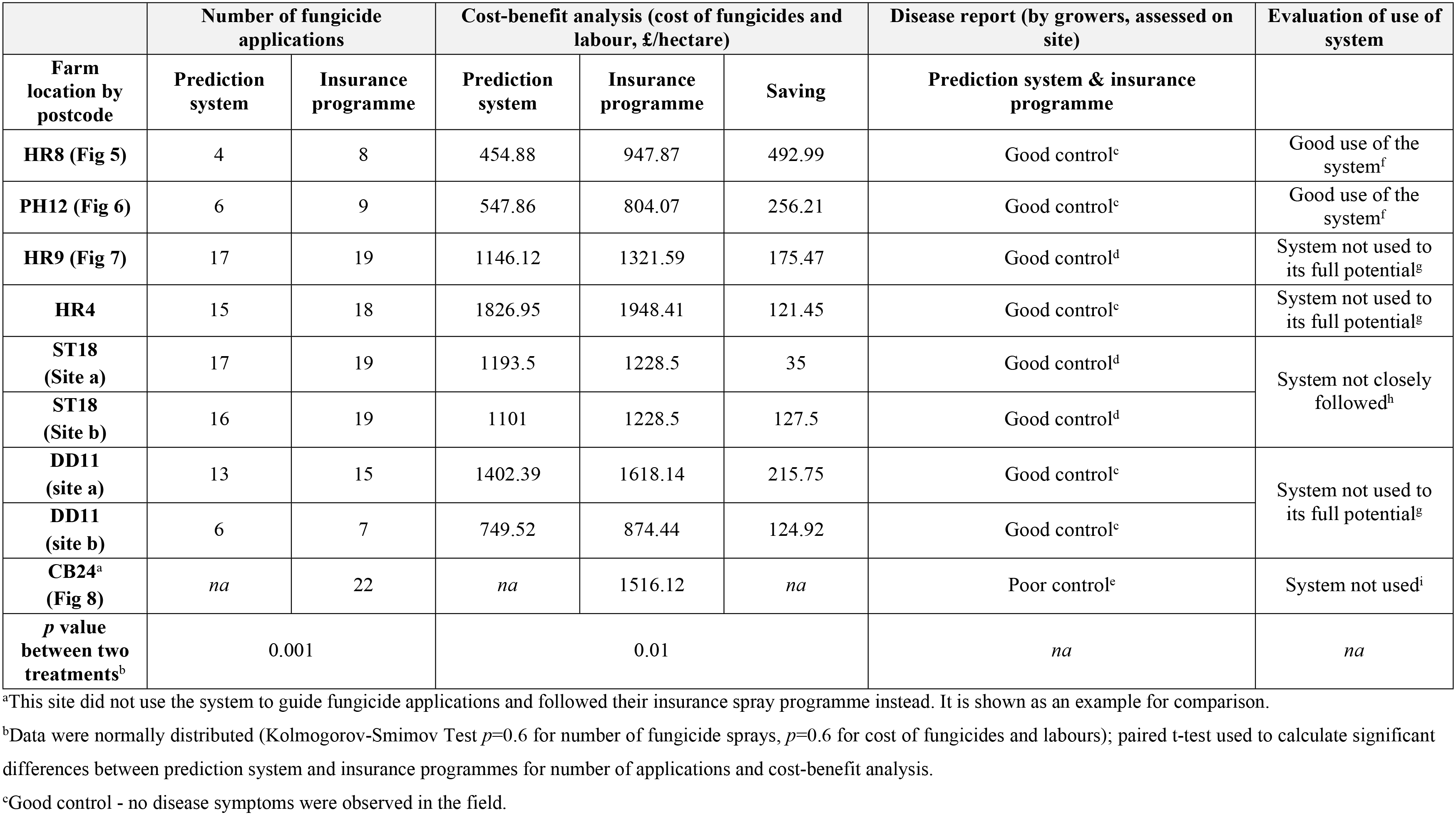
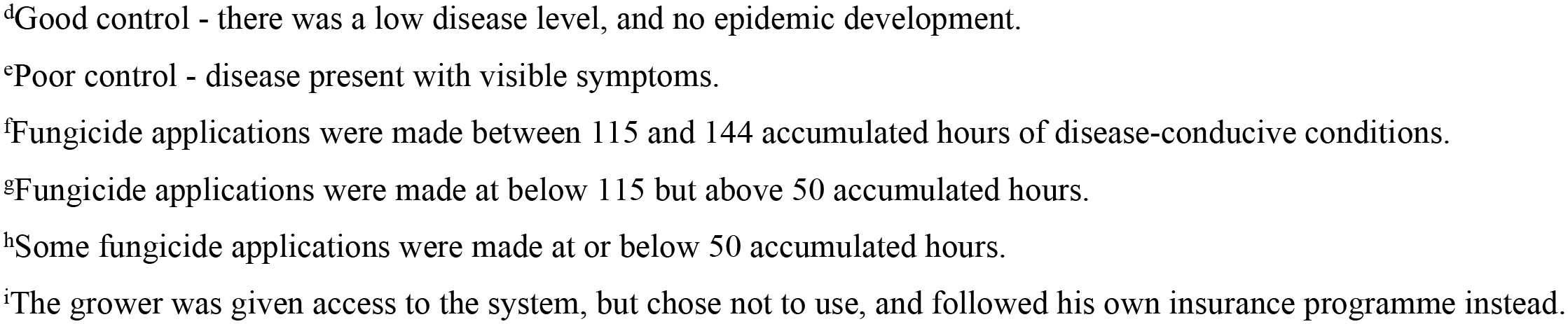
Results of on-farm validation on nine sites from seven farms (CB24 control site) in stage 3.

|  | Number of fungicide applications |  | Cost-benefit analysis (cost of fungicides and labour, £/hectare) |  |  | Disease report (by growers, assessed on site) | Evaluation of use of system |
| --- | --- | --- | --- | --- | --- | --- | --- |
| Farm location by postcode | Prediction system | Insurance programme | Prediction system | Insurance programme | Saving | Prediction system & insurance programme |  |
| HR8 (Fig 5) | 4 | 8 | 454.88 | 947.87 | 492.99 | Good control<sup>c</sup> | Good use of the system<sup>f</sup> |
| PH12 (Fig 6) | 6 | 9 | 547.86 | 804.07 | 256.21 | Good control<sup>c</sup> | Good use of the system<sup>f</sup> |
| HR9 (Fig 7) | 17 | 19 | 1146.12 | 1321.59 | 175.47 | Good control<sup>d</sup> | System not used to its full potential<sup>g</sup> |
| HR4 | 15 | 18 | 1826.95 | 1948.41 | 121.45 | Good control<sup>c</sup> | System not used to its full potential<sup>g</sup> |
| ST18 (Site a) | 17 | 19 | 1193.5 | 1228.5 | 35 | Good control<sup>d</sup> | System not closely followed<sup>h</sup> |
| ST18 (Site b) | 16 | 19 | 1101 | 1228.5 | 127.5 | Good control<sup>d</sup> |  |
| DD11 (site a) | 13 | 15 | 1402.39 | 1618.14 | 215.75 | Good control<sup>c</sup> | System not used to its full potential<sup>g</sup> |
| DD11 (site b) | 6 | 7 | 749.52 | 874.44 | 124.92 | Good control<sup>c</sup> |  |
| CB24<sup>a</sup> (Fig 8) | *na* | 22 | *na* | 1516.12 | *na* | Poor control<sup>e</sup> | System not used<sup>i</sup> |
| *p* value between two treatments<sup>b</sup> | 0.001 |  | 0.01 |  |  | *na* | *na* |

<sup>a</sup>This site did not use the system to guide fungicide applications and followed their insurance spray programme instead. It is shown as an example for comparison.
<sup>b</sup>Data were normally distributed (Kolmogorov-Smimov Test *p*=0.6 for number of fungicide sprays, *p*=0.6 for cost of fungicides and labours); paired t-test used to calculate significant differences between prediction system and insurance programmes for number of applications and cost-benefit analysis.
<sup>c</sup>Good control - no disease symptoms were observed in the field.
<sup>d</sup>Good control - there was a low disease level, and no epidemic development. <sup>e</sup>Poor control - disease present with visible symptoms. <sup>f</sup>Fungicide applications were made between 115 and 144 accumulated hours of disease-conducive conditions. <sup>g</sup>Fungicide applications were made at below 115 but above 50 accumulated hours. <sup>h</sup>Some fungicide applications were made at or below 50 accumulated hours. <sup>i</sup>The grower was given access to the system, but chose not to use, and followed his own insurance programme instead.

Readouts from the prediction system gave good indications of how growers used the system as well as their willingness and/or confidence to rely on its warnings to spray. Fig 5 shows that the best grower (site HR8) recorded all fungicide applications made, by resetting the system to zero (Fig 5, vertical lines) and three out of four sprays (Fig 5, Jul 20, Aug 5 and Aug 31) were performed between 115 and 144 accumulated hours (medium and high risk), while one fungicide (Jun 30) was applied at 80 accumulated hours (low risk). Using the prediction system extended the interval between the four fungicide applications by 20 days, 16 days and 26 days, respectively (Fig 5), thus reducing the number of applications made, compared to the insurance programme (7-14 days intervals).

**Fig 5.**
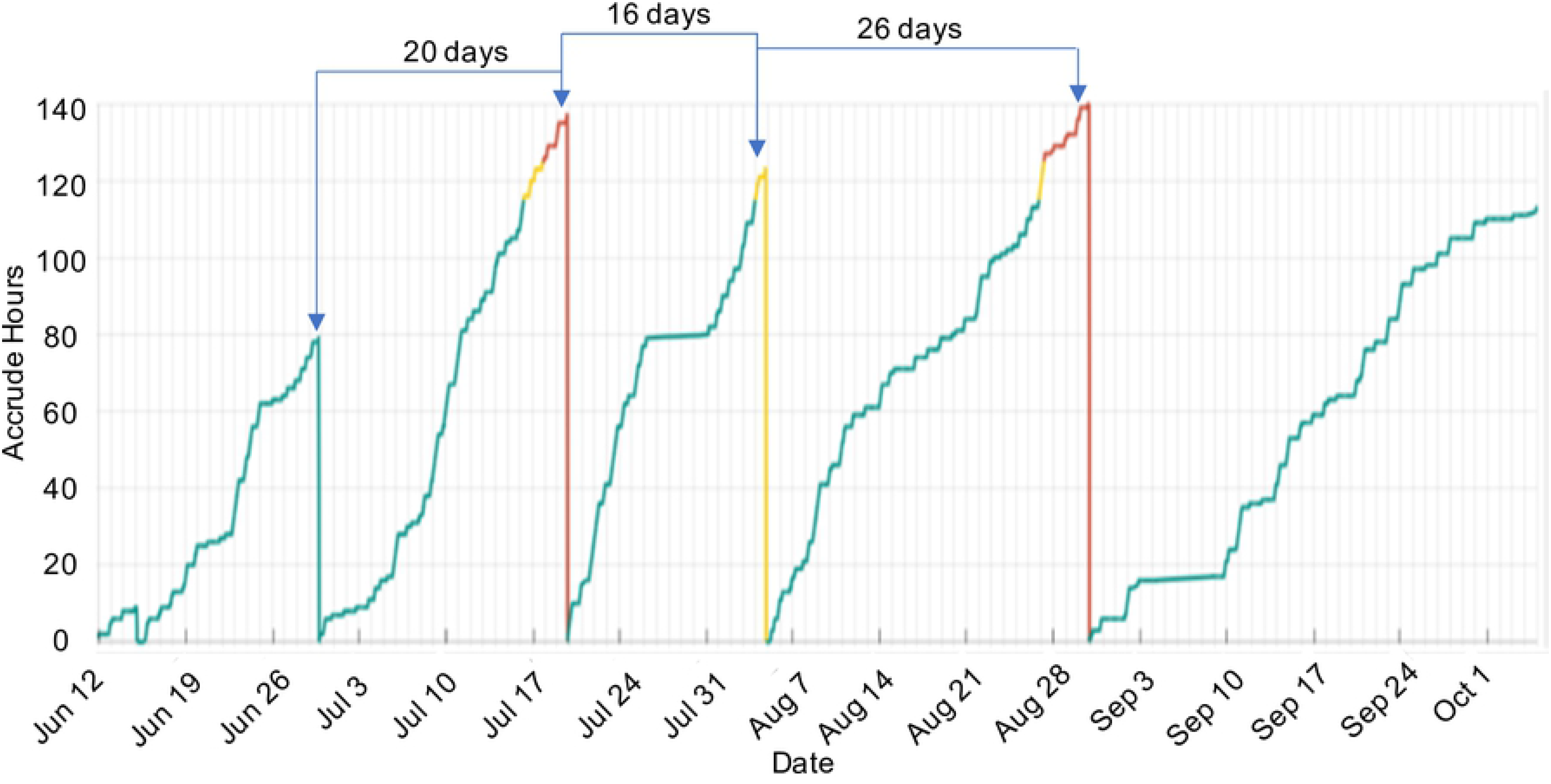
Screenshot of prediction graph at the end of season showing accumulation of disease-conducive conditions and applications of fungicides for site HR8 in 2019 (stage 3). This output of the prediction system utilised a ‘traffic light’ system to signify recommendations when growers should apply a fungicide. The Y-axis of the prediction graph indicates the number of accumulated hours where both parameters (temperature and RH) are met, the X-axis shows the date. When the grower logs into the real time web-based prediction system, the ascending line indicates the current status of the accumulated number of hours of suitable conditions, enabling the grower to decide whether a fungicide spray is required. When the ascending green line turns to amber (at 115 hours), the grower should prepare to spray. When the line turns to red (at 125 hours), a fungicide spray is needed. A fungicide should be applied before the elapsed time reaches 144 accumulated hours, to prevent *P. aphanis* sporulation. After each application, the fungicide details were entered by the grower and the system was reset. The number of days above the lines are the intervals between each spray. In this graph, the system has been reset four times; each time a fungicide application has been made, shown as a straight-line down to zero. The longest interval between two sprays was 26 days.

At site PH12, the grower used the predictions system reasonably well, but was not willing to allow the system to run beyond 115 accumulated hours before applying a fungicide (Fig 6). In this case, two fungicides (Fig 6, Jun 21 and Jul 10) were applied before 50 accumulated hours (low risk) and four fungicides (Fig 6, Jul 2, Jul 23, Aug 2 and Aug 15) were applied between 50 and 115 accumulated hours (low to medium risk). This grower managed to extend the interval between applications by three days throughout the season by using the prediction system, when compared to their insurance spray programme of 10-day intervals.

**Fig 6.**
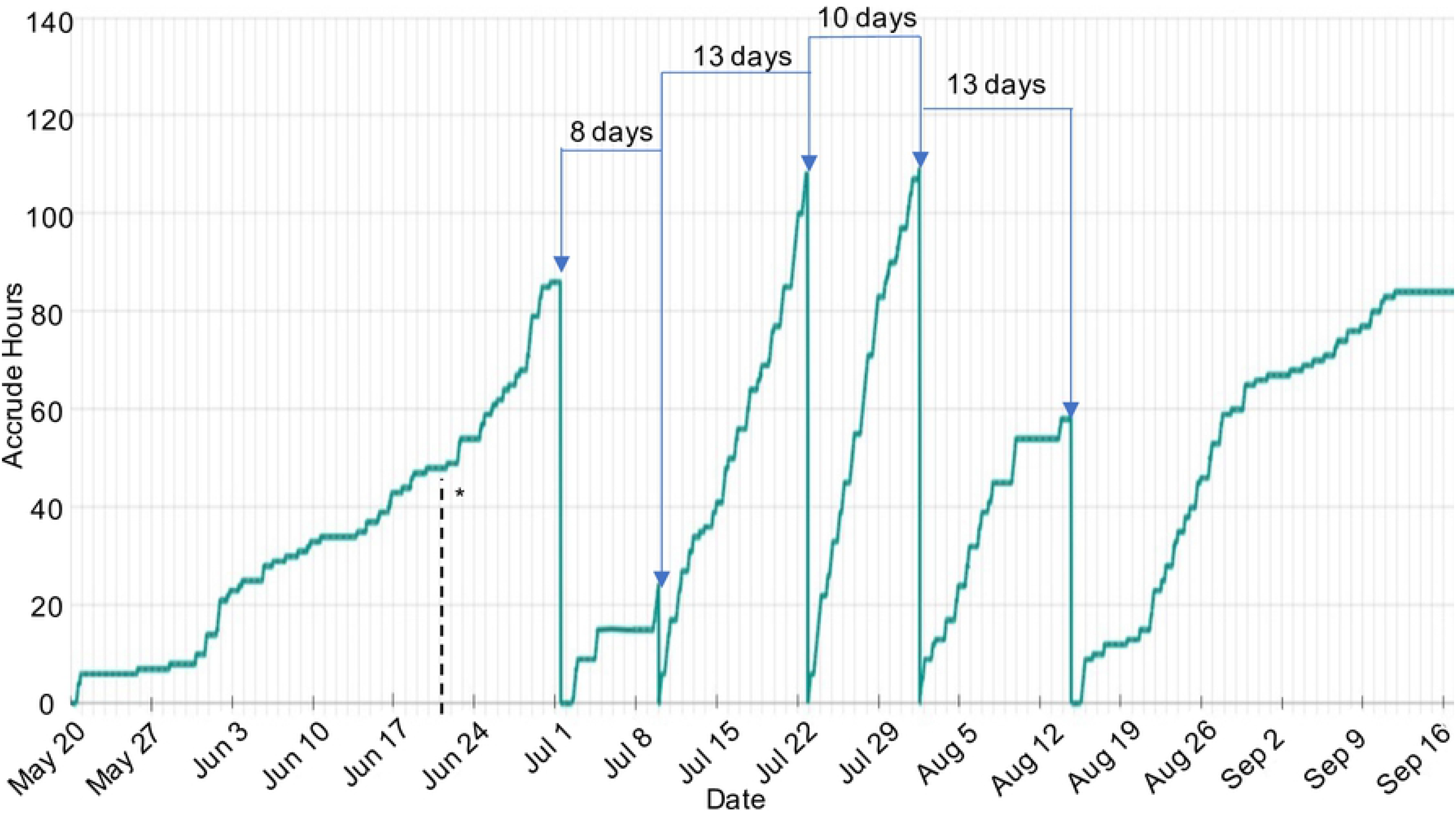
Screenshot of prediction graph at the end of season and applications of fungicides for site PH12 in 2019 (stage 3). In this graph, the system has been reset six times. The intervals between two sprays were between 8 and 13 days. ---* indicated that a fungicide was applied, but the prediction system was not reset.

In contrast, the grower from site HR9 was more erratic in willingness to rely on the prediction system for timing fungicide applications, and Fig 7 shows that the system was not followed strictly, resulting in a total of nine sprays (May 28, Jun 3, Jun 11, Jun 17, Sep 1, Sep 5, Sep 7, Sep 11 and Sep 17) being applied before 50 accumulated hours (low risk), mainly in the period from May 6 to June 24 and between Sept 2 to Sept 16. In between times, the grower allowed the system to exceed 115 accumulated hours (medium to high risk) on four occasions before applying a fungicide (Fig 7, Apr 26, Jul 15, Jul 29 and Aug 19) and the system was not reset to zero as it should have been after the application on Oct 7. Fig 9 shows that three fungicides (Jun 26, Jul 19 and Aug 27) applied between 50 and 115 hours (low to medium risk). A ‘clean up’ spray was applied on 26^th^ April, as required when using the prediction system. Using the prediction system increased the spray interval at the beginning of the season.

**Fig 7.**
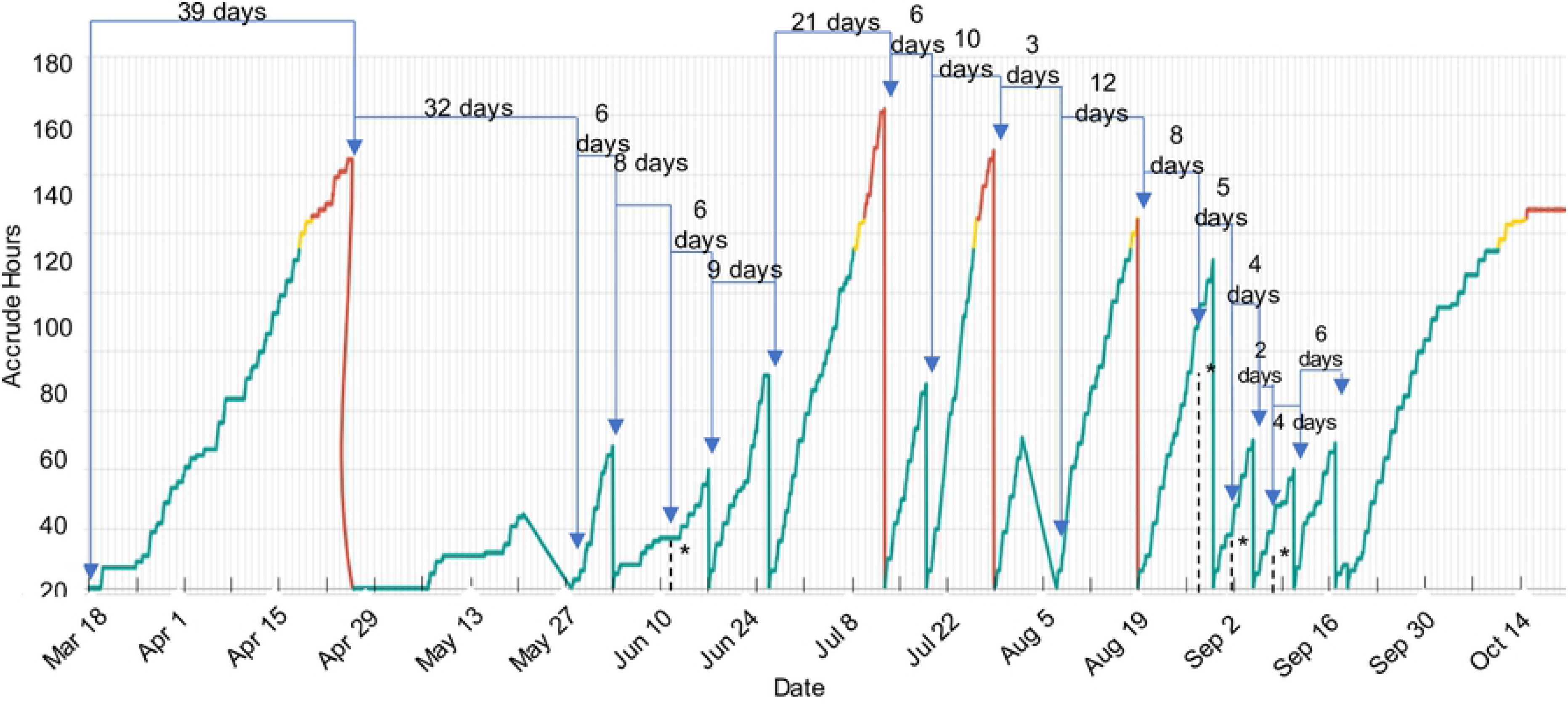
Screenshot of prediction graph at the end of season and applications of fungicides for site HR9 in 2019 (stage 3). In this graph, the system has been reset 17 times. The intervals between two sprays were longer (i.e. 39 days) at the start of the season and very short (i.e. <5 days) towards the end of the season. ---* indicated that a fungicide was applied, but the prediction system was not reset.

Fig 8 shows the recording of the insurance programme of frequent sprays (every five to seven days on average) at site CB24, a control farm where the prediction system was not used to guide fungicide applications. During the study, this grower accessed the prediction system from 21^st^ May 2019, before which three fungicide applications had already been made and were therefore not recorded in the readout. The grower did not use the prediction system to guide the timing of fungicide sprays, and Fig 8 shows that relying on frequent routine sprays resulted in many unnecessary fungicide treatments. Ten fungicide applications (Fig 8, May 22, May 25, Jun 1, Jun 6, Jun 7, Jun 15, Jun 19, Jun 22, Jul 1 and Jul 13) were made when the system was <50 accumulated hours (low risk) and one fungicide (Fig 8, Jun 27) was applied between 50 and 115 accumulated hours (low to medium risk).

**Fig 8.**
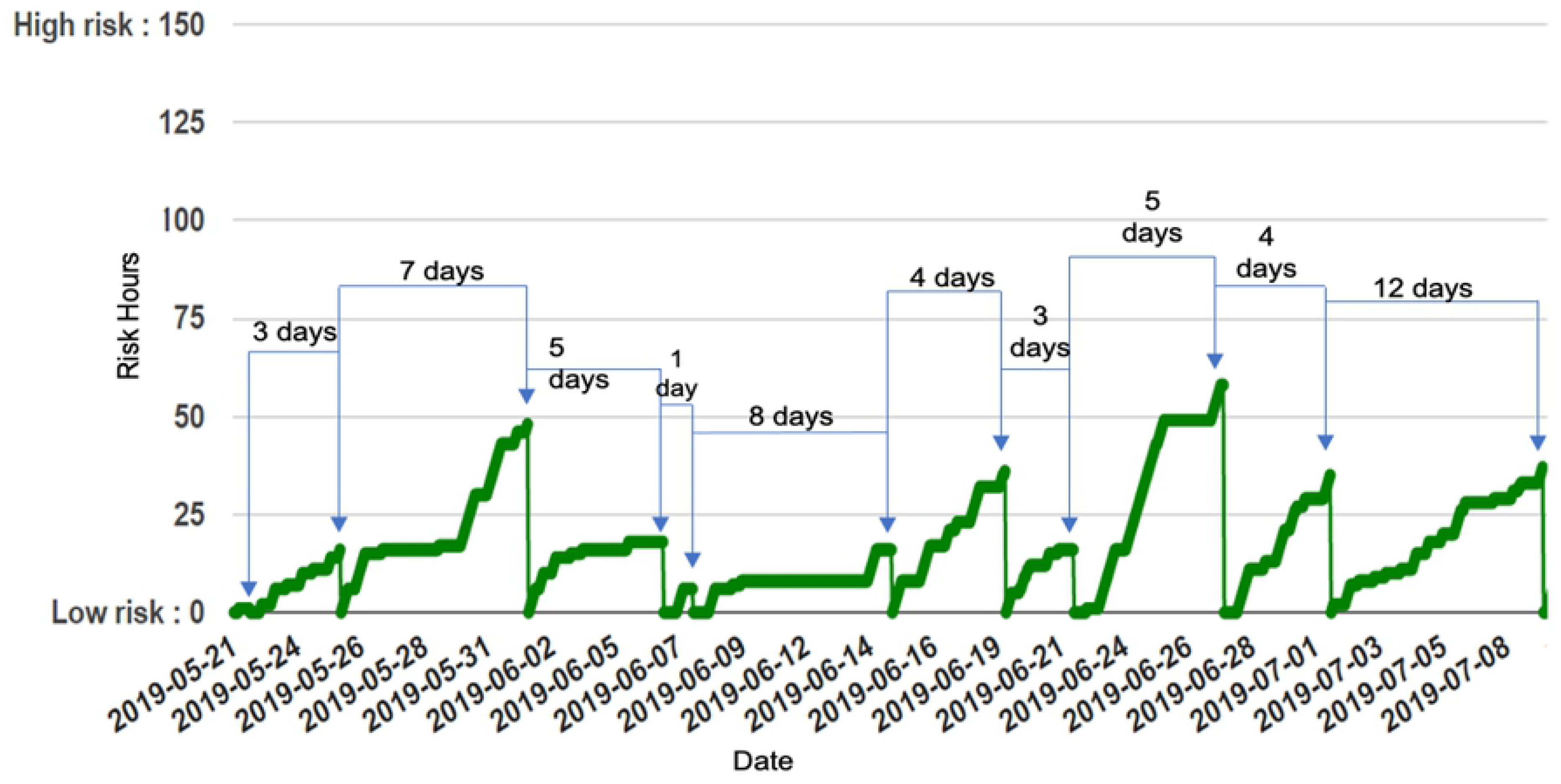
Screenshot of prediction graph at the end of season for CB24 (control site) that did not follow the prediction system in 2019 (stage 3). The grower followed his insurance spray programme and entered the fungicide details and reset the system after each spray. In this graph, 22 sprays were applied (three fungicide sprays were made prior to 21^st^ May 2019, but were not recorded in the system). Sprays were mostly made at <50 accumulated hours and the intervals between two sprays were between 1 and 12 days.

## Discussion

This paper has followed the development, over a 15-year period, of a rule-based strawberry powdery mildew risk prediction system that can be used as a decision support tool for determining the most effective timing for the application of fungicides. The system has been shown to have successfully enabled growers to reduce their fungicide use in strawberry crops, while maintaining good disease control. Good financial savings (between £35-£493/hectare) were achieved by the participating farms when using the system. Evidence from successful on-farm validations has shown that this real-time, user-friendly prediction system can be a very useful tool in the growers’ disease management programme.

Decision support tools for integrated pest/disease management are not new [30–34] and their success depends not only on their ability to monitor pests and pathogens, and to accurately predict action thresholds (i.e. minimum levels that justify pesticide treatment), but also on their ease of use and adoption by growers. The prediction system developed here is based on measuring the accumulated hours under pathogen conducive temperature (>15.5°C and <30°C) and relative humidity (>60%) conditions, which are accessible criteria and generally already recorded by growers for crop management. Unlike the recently published decision tree for forecasting SPMD [30], our system does not require the accurate determination of susceptible leaves or airborne inoculum levels as these were not essential for our system and might have limited the application of our system across a wider range of cultivars. Instead, it assumes the presence of disease inoculum and susceptibility of all crops, so that for the system to be effective, growers are advised to begin with a clean-up spray treatment (e.g. Fig 7, site HR9). Our previous research has shown that the initial inoculum can be quite high in over-wintered UK crops and that disease can also be imported from the propagators [15, 29]. The work described here was able to show that if the initial clean-up spray was neglected, as was the case with the grower at site CB24 (Fig 8), visible disease will occur, and the prediction system becomes ineffective for disease control. With correct use of the prediction system, as by the grower at site HR8 (Fig 5), we clearly demonstrated that epidemics of *P. aphanis* could be controlled with fewer fungicide applications.

Without the prediction system to guide them, or where growers were not willing to completely rely on the system, there was a tendency to overestimate how quickly *P. aphanis* was developing in their polythene tunnels, as they were concerned about the economic impact of severe disease, which caused them to use insurance sprays [25]. This behaviour was self-evident in the readout traces from the system (e.g. Fig 7, site HR9), indicating the application of fungicides when the system was showing a low risk of disease. This risk-averse attitude is also largely reflected in the higher frequency of fungicide applications recommended by insurance programmes for disease control.

To help growers gain confidence in using our system, and to manage the aversion to risk, we adopted a traffic light system to visually display the accumulation of disease-conducive hours with low (green), medium (amber) and high (red) risk periods clearly identified (e.g. Fig 5). This made it easy for growers to understand the data outputs and the practical implications. The use of the environmental parameters that had been developed and validated on commercial farms, rather than only in a laboratory, not only improved the precision of the system but also increased the confidence of growers in using it in commercial farm settings. The number of participating sites in the final validation (stage 3) increased from three in 2018 to nine in 2019 across England and Scotland. For those who followed the system well, good disease control was achieved. For example, site PE14 (Table 7) saw a lower disease incidence (a reduction of up to 45% in the AUDPC value) for both everbearer and June bearer crops in the prediction system plot compared with the untreated control. The expression of risk as disease-conducive hours enabled growers to understand that, for example, it was not necessary to frequently apply fungicides according to an insurance programme when only 50 out of 144 disease-conducive hours had been accumulated. In addition, the slope of the line in the graphical display output enabled the growers to follow the rate at which the disease risk was changing from medium to high risk, which allowed them to decide when to schedule a fungicide application and/or harvest window.

The prediction system has been tested on a range of cultivars (with different levels of disease susceptibility), with a selection of growing methods, at a range of geographical locations in both England and Scotland, using different types of weather sensors (Table 2), and proven to be effective in supporting growers to prevent epidemic build up and to control disease (Table 9). This indicates the robust usability of the prediction system. Furthermore, this also demonstrated an important feature that the disease epidemics generally develop in a similar manner wherever they occur. In other words, exposure to identical disease-conducive conditions results in the same level of disease development irrespective of the setting, whether it is in polythene tunnels or at geographical locations with various seasonal conditions.

Technological constraints were once recognized as one of the main obstacles for the widespread adoption of decision support systems in the 1990s [13, 35]. The situation has been much improved now thanks to the fast development of new technologies, which enables wider access to computers, internet and wi-fi services [35]. However, a much greater level of focus has recently been placed on the implementation of advanced technologies during the development of a DSS, with little thought given to the practicality of the system and the acceptance by its end-users i.e. growers [36–38]. In reality, many other factors can influence the acceptability of a DSS, such as user-friendliness, accessibility and complexity, etc. [36,39,40]. These should all be considered and be integrated within the decision-making process [36]. Initially (stage 1), encouraged by the increase in availability of personal computers on farms, we devised a process that required manual retrieval of relative humidity and temperature data from the data loggers and transfer to an Excel spreadsheet, to compute the disease risk. Growers found this to be too cumbersome and time-consuming. They needed to be able to access the data quickly, easily and frequently and at that time, this was addressed by using WeatherLink^TM^ enabled Davis weather stations for automated temperature and relative humidity data acquisition, and this was combined with the prediction system software on a local CD (stage 2). Unfortunately, the manual data download from the WeatherLink ^TM^ cloud computers and upload to the CD format did not sufficiently improve usability of the prediction system. The move to online real-time, wi-fi enabled technology in stage 3 rendered data acquisition and processing fully automatic and greatly improved the usability of the system. It enabled growers to quickly access the system online with relative ease and observe how disease-conducive hours were accumulating in real-time.

One of many factors that influences the adoption of a DSS is the growers’ willingness to take up new technology [11, 35]. It is recognised that participatory educational efforts, such as small class interactions, are necessary to engage a wider scale of growers; to discuss the scientific background of the work and to introduce and familiarise with the new software [13, 35]. It was found among growers that the perception of financial loss due to not applying insurance sprays can outweigh the cost savings that are potentially provided by the DSS [12, 41]. Therefore, in this work, grower confidence in the prediction system and its application was supported by education, delivered through knowledge exchange events. Plant pathology researchers and farm data acquisition and delivery technologists helped the growers to understand the technology as well as the biology of *P. aphanis* and pathogen infection and disease progression more clearly, which further increased their willingness to trust and adopt the prediction system to replace their insurance spray programmes. The provision of continuous support to growers through education e.g. seminars and workshops, etc. is believed to be essential to promote the long-term adoption of the DSS [11].

The innovative use of the real-time prediction system enabled growers to be more selective about the modes of action (MoA) of the fungicides applied. Growers have to consider modes of action used when making decisions about their spray programme [2]. There is only a limited number (<15) of active ingredients/groups that have been approved for use in strawberries with recommendations for powdery mildew control in the UK by 2019 [2]. Furthermore, *P. aphanis* has been found to have a medium to high level of insensitivity risk against eight out of the 14 fungicide MoA groups that are authorised for use on strawberries [42]. Therefore, products with the most effective anti-fungal active ingredients are generally reserved for periods at the end of the season when disease severity is greater, and fungicide applications are required more frequently. This deployment strategy, coupled with the fact that by using fewer sprays they did not exhaust the available MoA, gave the growers the opportunity to manage and mitigate the potential risk of fungicide insensitivity developing in the pathogen population.

The use of the prediction system not only improved the efficacy and precision of strawberry powdery mildew control but also saved associated fungicide and labour costs; average savings per hectare of £246 (over two sites) and £193.66 (over eight sites) were achieved in 2018 and 2019, respectively. When these values for savings are applied to the UK national crop area (4,774 ha in 2019) [1], an overall cost saving of between £924,533 and £1,174,404, equivalent to 0.3% of the national crop value, could be achieved. Thus, the financial savings for the individual grower are significant, and at the national level, it is the added environmental benefits accrued from reduced fungicide use that become more important. Studies have shown that the potential to demonstrate long-term economic, social and environmental sustainability is more important in encouraging the adoption of a DSS [11], than the short-term economic benefits alone [11, 43].

## Conclusion

The work reported here shows that a strawberry powdery mildew prediction system has been developed in the UK and works well on commercial farms. The system has been proven to be reliable, simple and easy to access. It serves as a decision support tool for informing decision making, and has been shown to be effective in guiding precise timings of fungicide application (i.e. with reduced fungicide applications compared to insurance sprays) leading to good commercially acceptable disease control. The simplicity of its criteria also makes it universally applicable to a wider range of strawberry cultivars and geographical locations. Moreover, successful disease control with fewer fungicide applications can be translated into tangible financial savings for growers. In all, this prediction system can meet the needs of the strawberry growers, enabling the production of disease-free crops, with fewer fungicide applications, with added benefits of cost savings, potentially fewer fungicide residues at harvest, as well as reduced risk of compromising fungicide MoA efficacy.

The prediction system was licenced and made commercially available in May 2020. Since then, a growing number of farms in the UK and abroad have either shown their interests or already taken on this product. This is encouraging as it shows not only growers’ willingness to embrace new technology (a result of continuous support through workshops and training sessions), and their real need for an effective decision support tool for their integrated farm management; more importantly, it demonstrates the great potential the prediction system can offer to assist them in that.

## Acknowledgements

We thank Henry Duncalfe, Harriet Duncalfe, Richard Hibbard, John Clark, David Dunn, Stuart Arbuckle and all participating growers in the 2019 validation for supplying field experimental sites and participating in the use of the prediction system. We thank KisanHub (Dr Sachin Shende) (2016-2018, Proof of concept) for the provision of a web-based prediction system; we also thank Agri-tech Services for collaboration in the 2019 on-farm validation. We thank Professor Xiangming Xu (NIAB-EMR) for his involvement in the DEFRA-AHDB Link project HL0191 (AHDB 2009). We also thank Terry Trinca and Eamonn Doherty, who took part in field experiments (2014 and 2015), and finally we thank Carmilla Asiana for her involvement in disease assessments (2017).

## Supporting information

**S1 Fig. Types of polythene tunnels for strawberry production in the UK.** (a) Spanish tunnel containing a strawberry crop. 1- raised bed, 2- strawberry crop planted in coir and 3- polythene tunnel; (b) Fleece covered strawberry crop in Spanish tunnel in spring. 1- coir bag containing young strawberry plants with irrigation drippers placed within bag, 2- fleece covering, 3- raised bed and 4- polythene tunnel; (c) Seaton tunnel containing strawberry crop. 1- polythene of Seaton tunnel, 2- permanent venting holes; (d) tabletop growing method. 1- tabletops, 2- strawberry crop grown in coir bags and 3- polythene tunnel.

**S2 Fig. Locations of experimental sites.** Key indicates locations of field experiments accompanied by first three or four digits of postcodes and dates experiments were undertaken.

## References

1. Defra. Horticultural Statistics 2019. [cited 2021 April 15]. Available from: https://www.gov.uk/government/statistics/latest-horticulture-statistics

2. Hall AM, Jin XL, Dodgson JLA. AHDB Factsheet 29/16: Control of strawberry powdery mildew under protection, Projects SF62, SF62a & SF113. 2016 [cited 2019 October 20]. Available from: https://ahdb.org.uk/knowledge-library/control-of-strawberry-powdery-mildew-under-protection-2

3. Peres NA, Mertely JC. Powdery mildew of strawberries. EDIS. 2013; 5. [cited 2021 April 29]. Available from: https://doi.org/10.32473/edis-pp129-2013

4. Looser N, Scherbaum E, Anastassiades M, Zipper H. Pesticide Residues in Strawberries sampled from the Market of the Federal State of Baden-Württemberg in the Period between 2002 and 2005. J Verbrauch Lebensm. 2006; 1, 135–141.

5. Wightwick A, Walters R, Allinson G, Reichman S, Menzies N. Environmental risks of fungicides used in horticultural production systems. In Carisse O, editor. Fungicides. London: IntechOpen; 2010. pp.273–304.

6. Bourke PA. Use of weather information in the prediction of plant disease epiphytotics. Annu. Rev. Phytopathol. 1970; 8, 345–370.

7. Gent DH, Mahaffee WF, McRoberts N, Pfender WF. The use and role of predictive systems in disease management. Annu. Rev. Phytopathol. 2013; 51, 267–289.

8. Hughes G. The evidential basis of decision making in plant disease management. Annu. Rev. Phytopathol. 2017; 55, 41–59.

9. Jørgensen LN, van den Bosch F, Oliver RP, Heick TM, Paveley ND. Targeting fungicide inputs according to need. Annu. Rev. Phytopathol. 2017; 55, 181–203.

10. Taylor MC, Hardwick NV, Bradshaw NJ, Hall AM. Relative performance of five forecasting schemes for potato late blight (*Phytophthora infestans*) I. Accuracy of infection warnings and reduction of unnecessary, theoretical, fungicide applications. Crop Prot. 2003; 22, 275–283.

11. Rossi V, Sperandio G, Caffi T, Simonetto A, Gilioli G. Critical Success Factors for the Adoption of Decision Tools in IPM. J. Agron. 2019; 9, 710. 38. Wearing CH. Evaluating the IPM implementation process. Annu. Rev. Entomol. 1988; 33, 17–38.

12. Rossi V, Caffi T, Salinari F. Helping farmers face the increasing complexity of decision-making for crop protection. Phytopathol Mediterr. 2012: 51, 457–479.

13. Gent DH, DeWolf E, Pethybridge SJ. Perceptions of risk, risk aversion, and barriers to adoption of decision support systems and integrated pest management: an introduction. Phytopathology. 2010; 101, 640–643.

14. Xu XM, Robinson J, Simpson D. Interactions between isolates of powdery mildew (*Podosphaera aphanis*) and cultivars of strawberry, *Fragaria x ananassa*. Integrated Plant Protection for Soft Fruit. IOBC/WPRS Bull. 2008; 39, 203–209.

15. Jin XL. Epidemiology and control of powdery mildew (*Podosphaera aphanis*) on strawberry, PhD Thesis, University of Hertfordshire. 2016. Available from: https://uhra.herts.ac.uk/handle/2299/17212.

16. Amsalem L, Freeman S, Rav-David D, Nitzani Y, Sztejnberg A, Pertot I et al. Effect of climatic factors on powdery mildew caused by Sphaerotheca macularis f. sp. fragariae on strawberry. Eur. J. Plant Pathol. 2006; 114, 283–292.

17. Blanco C, De Ios Santos B, Barrau C, Arroyo F, Porras M, Romero F. Relationship among concentrations of *Sphaerotheca macularis* conidia in the air, environmental conditions, and the incidence of powdery mildew in strawberry. Plant Dis. 2004; 88, 878–881.

18. Jhooty JS, McKeen WE. The influence of host leaves on germination of the asexual spores of *Sphaerotheca macularis*. Can. J. Microbiol. 1965; 11, 539–545.

19. Jhooty JS, McKeen WE. Studies on powdery mildew of strawberry caused by *Sphaerotheca macularis*. Phytopathology. 1965; 55, 281–285.

20. Miller TC, Gubler WD, Geng S, Rizzo DM. Effects of temperature and water vapor pressure on conidial germination and lesion expansion of *Sphaerotheca macularis* f. sp. *fragariae*. Plant Dis. 2003; 87, 484–492.

21. Peries OS. Studies on strawberry mildew, caused by *Sphaerotheca macularis* (Wallr. ex Fries) Jaczewski I. Biology of the fungus. Ann. Appl. Biol. 1962a; 50, 211–224.

22. Peries OS. Studies on strawberry mildew, caused by *Sphaerotheca macularis* (Wallr. ex Fries) Jaczewski. II. Host-parasite relationships on foliage of strawberry varieties. Ann. Appl. Biol. 1962b; 50, 225–233.

23. Berger RD, Hau B, Weber GE, Bacchi LMA, Bergamin Filho A, Amorim L. A simulation model to describe epidemics of rust of phaseolus beans I. Development of the model and sensitivity analysis. Phytopathology. 1995; 85, 715–721.

24. Willocquet L, Savary S. An epidemiological simulation model with three scales of spatial hierarchy. Phytopathology. 2004; 94, 883–891.

25. Wise K, Mueller D. Are fungicides no longer just for fungi? An analysis of foliar fungicide use in corn. APSnet Features. 2011; Available from: https://www.apsnet.org/edcenter/apsnetfeatures/Pages/fungicide.aspx. doi:10.1094/APSnetFeature-2011-0531

26. AHDB. Grower Summary: SF 094/ HL0191: Minimising pesticide residues in strawberry through integrated pest, disease and environmental crop management (HortLINK). 2009 [cited 2019 September 05]. Available from: https://ahdb.org.uk/sf-094-minimising-pesticide-residues-in-strawberry-through-integrated-pest-disease-and-environmental-crop-management-hortlink

27. The Kolmogorove-Smirnov Test of Normality [Internet]. c2021 [cited 2021 Jun 28]. Social Science Statistics. Available from: https://www.socscistatistics.com/tests/kolmogorov/default.aspx

28. Dodgson JLA. Epidemiology and sustainable control of *Podosphaera aphanis* (strawberry powdery mildew), PhD Thesis, University of Hertfordshire. 2007. Available from: https://uhra.herts.ac.uk/handle/2299/14356

29. Liu B. Sustainable strawberry production and management including control of strawberry powdery mildew, PhD Thesis, University of Hertfordshire. 2017. Available from: https://uhra.herts.ac.uk/handle/2299/19051

30. Carisse O. Fall ML. Decision Trees to Forecast Risks of Strawberry Powdery Mildew Caused by *Podosphaera aphanis*. Agriculture. 2021; 11, 29. Available from: https://doi.org/10.3390/agriculture11010029

31. Carisse OL. Van der Heyden H. A new risk indicator for botrytis leaf blight of onion caused by Botrytis squamosa based on infection efficiency of airborne inoculum. Plant Pathol. 2012; 61, 1154– 1164.

32. Eccel E, Fratton S, Ghielmi L, Tizianel A, Shtienberg D, Pertot I. Application of a non-linear temperature forecast post-processing technique for the optimization of powdery mildew protection on strawberry. Ital. J. Agrometeorol. 2010; 2, 5–14.

33. Bardet A. Vibert J L’oïdium du fraisier. Un outil de prévision du risque. Infos Ctifl. 2011; 276, 38– 44.

34. Hoffman LE. Gubler WD. Validation of the UC Davis Strawberry Powdery Mildew Risk Index. Calif. Strawb. Comm. Rep. 2002; 2, 19.

35. Jones VP, Brunner, JF, Grove GG, Petit B, Tangren GV, Jones WE. A web-based decision support system to enhance IPM programs in Washington tree fruit. Pest Manag. Sci. 2010; 66, 587–595.

36. Matthews KB, Schwarz G, Buchan K, Rivington M, Miller D. Wither agricultural DSS? Comput. Electron. Agric. 2008; 61, 149–159.

37. Van Delden H, Luja P, Engelen G. Integration of multi-scale dynamic spatial models of socio-economic and physical processes for river basin management. Environ. Modell. Softw. 2007; 22, 223–238.

38. Worm GIM, van der Helm AWC, Lapikas T, van schagen KM, Rietveld LC. Integration of models, data management interfaces and training support in a drinking water treatment plant simulator. Environ. Modell. Softw. 2010; 25, 677–683.

39. Magarey RD, Travis JW, Russo JM, Seem RC, Magarey PA. Decision Support Systems: Quenching the Thirst. Plant Dis. 2002; 86, 4–14.

40. Kerr D. Factors influencing the development and adoption of knowledge based decision support systems for small, owner-operated rural business. Artif. Intell. Rev. 2004; 22, 127–147.

41. Wearing CH. Evaluating the IPM implementation process. Annu. Rev. Entomol. 1988; 33, 17–38.

42. FRAG. Fungicide resistance management in soft fruit. 2019. [cited 2021 Jun 10]. Available from: https://ahdb.org.uk/frag

43. Oliver DM, Fish RD, Winter M, Hodgson CJ, Heathwaite AL, Chadwick DR. Valuing local knowledge as a source of expert data: Farmer engagement and the design of decision support systems. Environ. Model. Softw. 2012; 36, 76–85

